# Heterologous iron-sulfur cluster biogenesis and delivery for cytosolic isobutanol and isopentanol production in *Saccharomyces cerevisiae*

**DOI:** 10.64898/2026.05.29.728687

**Authors:** Jeremy D. Cortez, José L. Avalos

**Affiliations:** Department of Molecular Biology;, Princeton University, Princeton, NJ; Department of Chemical and Biological Engineering;, Princeton University, Princeton, NJ; Andlinger Center for Energy and the Environment;, Princeton University, Princeton, NJ; Omenn-Darling Bioengineering Institute, Princeton University, Princeton, NJ

**Keywords:** *Iron-sulfur clusters*, *cytosolic pathway*, *bioinformatics*, *branched-chain higher alcohols*, *isobutanol*, *isopentanol*, *Saccharomyces cerevisiae*

## Abstract

*Saccharomyces cerevisiae* is an excellent microbial platform for sustainable production of next generation biofuels such as the branched chain higher alcohols (BCHAs) isobutanol and isopentanol. A cytosolic pathway for BCHA production is generated from expression of prokaryotic orthologs of branched-chain amino acid (BCAA) enzymes acetolactate synthase (ALS), mutant NADH-dependent ketol-acid reductoisomerase (KARI^P2D1-A1^), and dihydroxy-acid dehydratase (DHAD). The potential for this pathway has been hindered by the availability of iron-sulfur clusters, particularly the 2Fe-2S cluster, required for DHAD to function in the cytosol. *ILV3*, the endogenous yeast DHAD located in the mitochondria, can be deleted to create a valine auxotroph. In this study we use bioinformatics, heterologous gene library synthesis, and a valine complementation assay to find prokaryotic iron-sulfur cluster biosynthetic gene clusters (BGC) and accessory genes that aid DHAD function in the yeast cytosol. This work presents, to our knowledge, the first functional BGC that enhances the cytosolic activity of prokaryotic DHADs in *S. cerevisiae*. The SUF BGC from *Bacillus subtilis* combined with a ferritin-like protein (FTNB) from *Escherichia coli* and the *Lactococcus lactis* DHAD enhanced the production of BCHAs. Combined expression gave an average isobutanol titer of 412mg/L, 1.8-fold greater than *L. lactis* DHAD expressed alone. This work establishes a blueprint for better biofuel production by improving iron-sulfur cluster dependent enzyme activity in the yeast cytosol.

## 1 Introduction

To mitigate the environmental impact of fossil fuels, advanced biofuel alternatives must be produced more affordably and at larger scales (Fulton et al., 2015). Branched chain higher alcohols (BCHAs), such as isobutanol, isopentanol, and 2-methyl-1-butanol, are considered next generation biofuels because of their advantageous fuel properties (Choi et al., 2014). Compared to ethanol, BCHAs are a better option for ignition engines due to their lower vapor pressure, higher energy density, and better compatibility with current gasoline infrastructure (Fu et al., 2021). *Saccharomyces cerevisiae* naturally produces BCHAs via the Ehrlich degradation pathway of branched chain amino acids (BCAAs) valine, isoleucine, and leucine. While natural production is small, many metabolic engineering efforts have significantly increased BCHA production compared to natural levels (Gambacorta et al., 2022).

As depicted in Fig. 1A, *S. cerevisiae* converts glucose to pyruvate via glycolysis, which can be further metabolized to produce ethanol, BCAAs, and BCHAs (Kohlhaw, 2003). Some of the pyruvate is shuttled into the mitochondria via the mitochondrial pyruvate transport complex, where two pyruvate molecules are condensed by acetolactate synthase (ALS, encoded by *ILV2*). Subsequently, a nicotinamide adenine dinucleotide phosphate (NADPH)-dependent ketol-acid reductoisomerase (KARI, encoded by *ILV5*) converts the acetolactate to 2,3-dihydroxy isovalerate (Park and Hahn, 2019). Also in mitochondria, the 2Fe-2S-dependent dihydroxy-acid dehydratase (DHAD, encoded by *ILV3*) catalyzes formation of α-ketoisovalerate (α-KIV) from 2,3-dihydroxy-isovalerate. In both the mitochondria and cytosol, α-KIV is acetylated to α-isopropylmalate (α-IPM) via isopropylmalate synthases encoded by *LEU4* and *LEU9*, constituting the committed step to leucine biosynthesis. Alternatively, α-KIV can either be transaminated by BCAA aminotransferases encoded by *BAT1* and *BAT2* to produce valine or decarboxylated and reduced by various α-ketoacid decarboxylases (KDCs) and alcohol dehydrogenases (ADHs), via the Ehrlich pathway, to produce isobutanol. In the cytosol, α-IPM can also be converted to α-ketoisocaproate (α-KIC) through sequential reactions catalyzed by isopropylmalate isomerase and b-isopropylmalate dehydrogenase (encoded by *LEU1* and *LEU2*, respectively). This α-KIC can be transaminated to leucine by bat2p, or degraded to isopentanol by KDCs and ADHs, similar to the α-KIV conversion to isobutanol. In an alternative pathway, if ilv2p condenses one pyruvate with one α-ketobutyrate (usually derived from threonine), it commits the pathway to isoleucine biosynthesis, in which the sequential activities of ilv5p and ilv3p produce α-keto-3-methylvalerate (α-K3MV). α-K3MV, similarly, can be transaminated to isoleucine by bat1p/bat2p, or degraded to 2-methyl-1-butanol by KDCs and ADHs (Kohlhaw, 2003). Ilv2p, ilv5p, ilv3p, bat1p, leu9p and a fraction of leu4p are compartmentalized in mitochondria, whereas the rest of leu4p, leu1p, leu2p, bat2p, and the Ehrlich pathway (KDCs, and ADHs) are localized in the cytosol (Fig. 1A).

**Figure 1.**
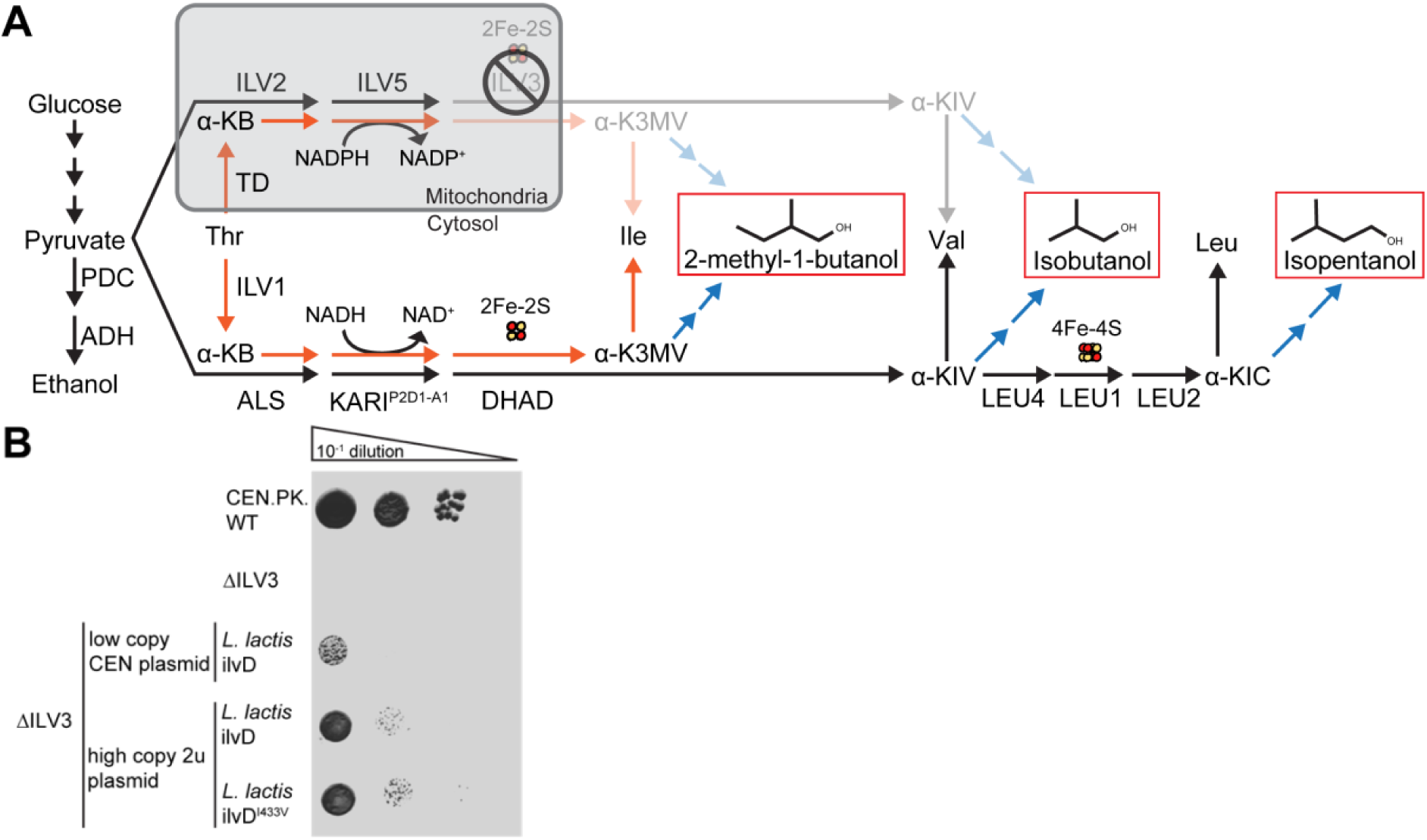
**Design and verification of valine auxotrophy**: (A) BCAA/BCHA pathway. Biosynthesis of branched-chain amino acids is parallel to the production of 2-methyl-1-butanol, isobutanol, and isopentanol (BCHAs outlined in red boxes). Isoleucine biosynthesis from threonine is denoted with the orange arrows. Valine and leucine biosynthesis from pyruvate is denoted by the black arrows. Parts of the pathway affected by *ILV3* deletion are made translucent. Mitochondrial parts of the pathway are shaded by the gray box. PDC, pyruvate decarboxylase; ADH, alcohol dehydrogenase; TD, threonine deaminase; ALS, acetolactate synthase; KARI^P2D1-A1^, NADH-dependent mutant of ketol-acid reductoisomerase; DHAD, 2Fe-2S dependent dihydroxy-acid dehydratase; α-KB, α-ketobutyrate; α-K3MV, α-keto-3-methylvalerate; α-KIV, α-ketoisovalerate; α-KIC, α-ketoisocaproate. (B) DHAD expressed in low and high copy plasmids rescue growth after an ILV3 deletion in synthetic complete media without valine.

The fractionation of BCAA biosynthesis and the Ehrlich pathway between mitochondria and cytosol has posed significant challenges in the production of BCHAs. Previous studies have co-localized these pathways to either the cytosol or mitochondria by removing or adding mitochondrial localization sequences (MLSs) to key enzymes (Brat et al., 2012; Avalos et al., 2013). Compartmentalizing the pathway in mitochondria improves production by increasing the local availability of enzymes, substrates, and co-factors, particularly 2Fe-2S clusters which are synthesized in this organelle (Lill and Freibert, 2020). However, the mitochondrial membranes pose additional challenges to the exchange of NAD^+^/NADH cofactors within the cytosol, which is required to approach theoretical yields (Gambacorta et al., 2022).

DHADs (such as ilv3p in yeast) require precious iron-sulfur cluster cofactors (2Fe-2S) mostly limited to the mitochondria, resulting in a major bottleneck for cytosolic isobutanol production (Gambacorta et al., 2022). Previous studies show that expression of a DHAD ortholog (ilvD from *Lactococcus lactis*) in the cytosol works for cytosolic isobutanol production but may be limited due to 2Fe-2S availability (Zhang et al., 2022). Overexpression of the yeast transcription factor *AFT1*, involved in iron homeostasis, or deletion of *FRA2*, which inhibits Aft1p nuclear translocation (Martinez-Pastor et al., 2017), increases isobutanol production likely because of increased iron-sulfur cluster availability (Gambacorta et al., 2022; Zhang et al., 2022). It is unknown whether expression of biosynthetic gene clusters (BGC) and auxiliary proteins that aid 2Fe-2S production in the yeast cytosol could further enhance isobutanol production.

Previous studies suggest that prokaryotic iron-sulfur cluster BGCs could be active when co-expressed with the correct accessory genes in the yeast cytosol (Biz and Mahadevan, 2021). For example, Carlsen et al. indicate that overexpressing components of the ISC BGC from *E. coli* alone does not produce a functional iron-sulfur cluster-dependent (E)-4-hydroxy-2-methylbut-2-enyl-diphosphate (HMBPP) synthase (ispG) of the methylerythritol phosphate (MEP) pathway (Carlsen et al., 2013; Partow et al., 2012). Only the co-expression of the ISC BGC with flavodoxin partners (fldA from *E. coli* and *B. subtilis*) results in an active HMBPP synthase and MEP pathway (Kirby et al., 2016). Co-expression of the correct iron-sulfur cluster BGC and accessory genes is challenging as the pathways and functions of certain iron-sulfur cluster related genes are still under investigation (Garcia et al., 2022). However, with an efficient high-throughput process and library, the correct genes to co-express could be found. To our knowledge, these co-expression attempts have yet to be reported for DHAD function and isobutanol production in yeast (Biz and Mahadevan, 2021).

In this study we utilize a cytosolic BCHA pathway previously found to be functional but limited by iron-sulfur cluster availability (Zhang et al., 2022). Our BCHA pathway combines ALS from *Bacillus subtilis* (Bs_AlsS), mutant NADH-dependent KARI from *E. coli* (Ec_ilvC^P2D1-A1^), and DHADs from multiple prokaryotic sources. We propose that, in addition to beneficial overexpression of *AFT1* (Zhang et al, 2022), heterologous iron-sulfur cluster BGC and accessory gene co-expression with the correct DHAD ortholog can improve the iron-sulfur cluster bottleneck. Because BCHAs are linked to BCAA production, this study utilizes a BCAA auxotroph with deleted endogenous DHAD (*ILV3*) and screens a heterologous library of DHAD enzymes and prospective iron-sulfur cluster related genes selected via a bioinformatic pipeline.

## 2 Methods

### 2.1 Bioinformatics and phylogenetic analysis

DHAD orthologs were chosen from a protein database generated from a PFAM hidden Markov model (HMM) motif (ilvDEDD, PF00920, PF24877) search of the Uniprot database (v2022.02) representing all known prokaryotic proteomes. We used *hmmsearch* from the HMMER v3.2.1 package for hits with E-value lower than 10^-4^ (McClure et al., 1996). This database was then filtered to combine any redundant sequences or serotypes under organisms defined by NCBI taxonomy. A final protein database of 371 members representing at least 1 genome from 374 bacteria (Supp. File 1), and the yeast sequence of *ILV3*, were aligned using MAFFT v7.526 and trimmed with BMGE v1.1 using the substitution matrix BLOSUM62. Maximum likelihood (ML) trees were generated using Fast-Tree v2.1.11 with the Jones-Taylor-Thorton model and viewed with the interactive tree of life (iTOL) display and annotation tool (Letunic and Bork, 2021; Price et al., 2010). Best representative orthologs to synthesize were selected from a consolidated ML-tree pruned to node resolution of at least 1 via Treemmer v0.3 (Menardo et al., 2018). Patristic distance to the yeast *ILV3* was generated from the unpruned ML-tree using the Bio.Phylo.BaseTree module from Biopython (Cock et al., 2009). The full bioinformatic pipeline and python script is available at https://github.com/jdcortez93/Avalos_bioinformatics.

Representative *E. coli* ISC and *B. subtilis* SUF BGC genes were selected for synthesis using BioCyc (biocyc.org) and a curated literature search. *L. lactis* SUF genes were identified by BLAST search of the *L. lactis* proteome with SUF related genes from *B. subtilis* (E-value results represented in Supp. Table 3). The most significant genes found (listed in Supp. Table 3) were codon optimized and synthesized with Twist Biosciences.

### 2.2 Genomic Neighborhood Network (GNN) mapping and neighbor ranking

For the initial proposed ortholog library, genomic coordinates of 100 prospective-DHAD neighboring genes upstream and downstream were gathered via UniProt ID lookup (v2022.02) of the Enzyme Function Initiative (EFI) database (Zallot et al., 2019). A python script was written to rank genes upstream and downstream of DHAD based on closest genomic distance. This ranking was used to select proposed DHAD orthologs with at least 1 KARI located 3 genes upstream or downstream (GNN3). Python script can be found at https://github.com/jdcortez93/Avalos_bioinformatics.

### 2.3 Plasmid library construction

Vectors for ortholog DHAD and iron-sulfur cluster accessory genes were generated with restriction enzyme and ligation cloning in collaboration with the Joint Genome Institute (JGI) using the pESC 2-micron plasmid as a parent. All heterologous genes were codon optimized for yeast with the JGI BOOST (v2) software tool and synthesized with Twist Biosciences. Each DHAD ortholog was paired with each one of iron-sulfur cluster accessory genes (selected via a curated literature search, Table 1) that could assist activity on a dual GAL promoter. The DHAD synthesized genes were assembled with BamHI digested pESC via Gibson assembly (Gibson et al., 2009). Each assembled DHAD plasmid was digested with NotI and assembled with iron-sulfur cluster accessory genes also via Gibson assembly. Gibson assemblies were transformed in *Escherichia coli* strain DH5α, selected, and arrayed in 96 well format with 30% glycerol for frozen stocks. Sequences were verified by Sanger sequencing. Plasmids were miniprepped after the arrayed 96 well cells were grown individually in LB with ampicillin. Arrayed cells grown overnight were pooled and plasmids purified with the MAX miniprep kit from Qiagen. All other plasmids in this study are described in Supp. Table 2. BGC plasmids were cloned via homologous recombination by transforming *S. cerevisiae* with plasmid parts containing 25-50 bp homology regions onto a CEN plasmid (p414). Each BGC gene was cloned with strong constitutive promoters and terminators known to function in yeast (Supp. Table 2).

**Table 1.**
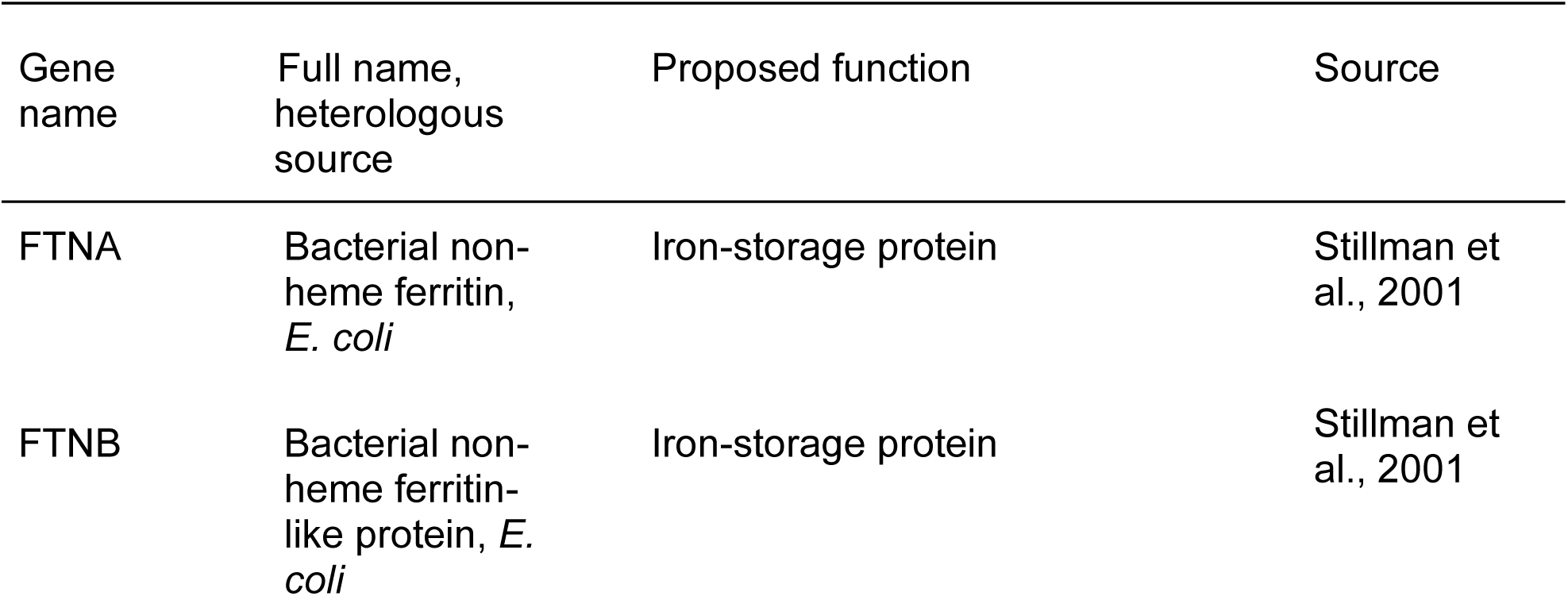
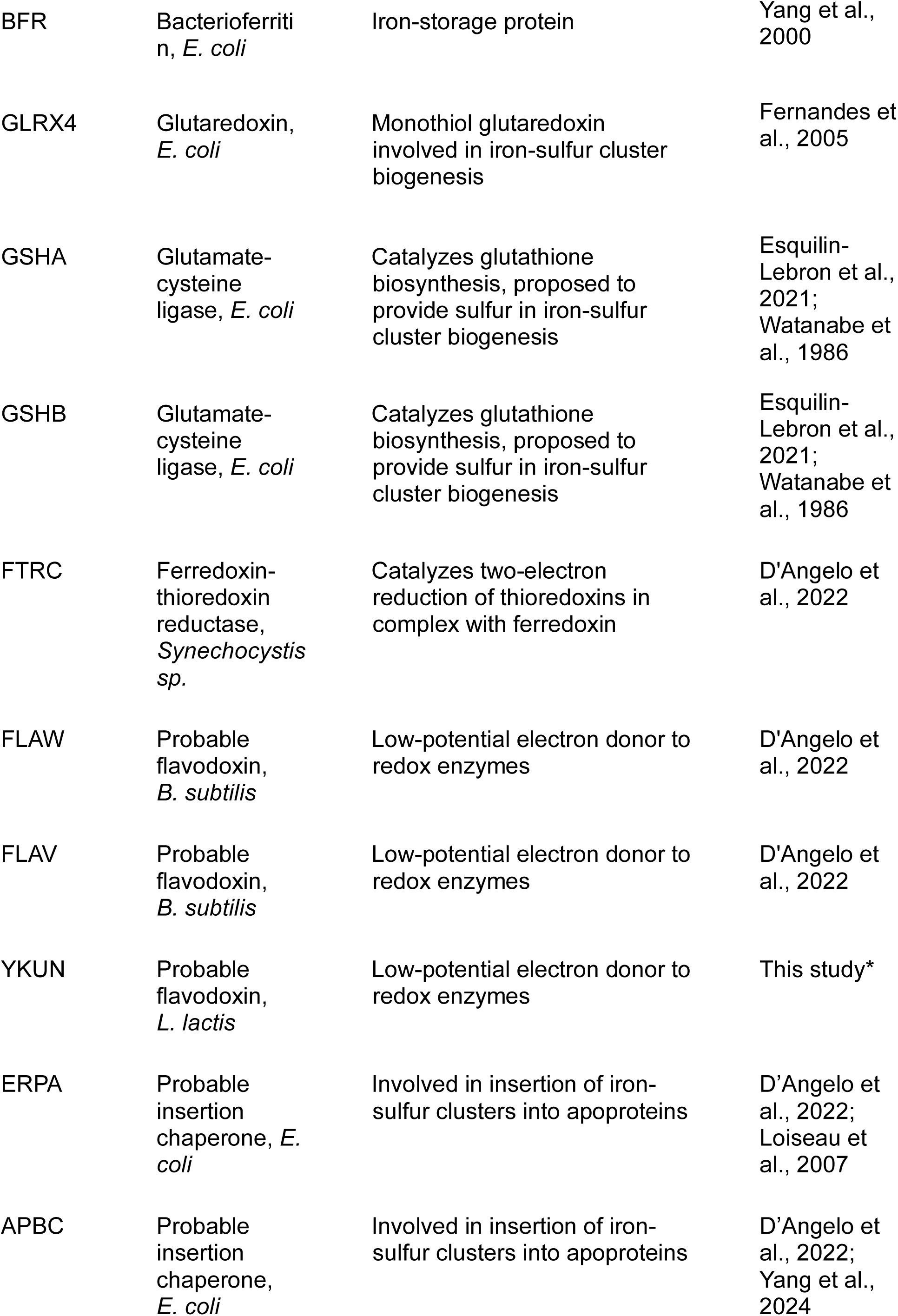
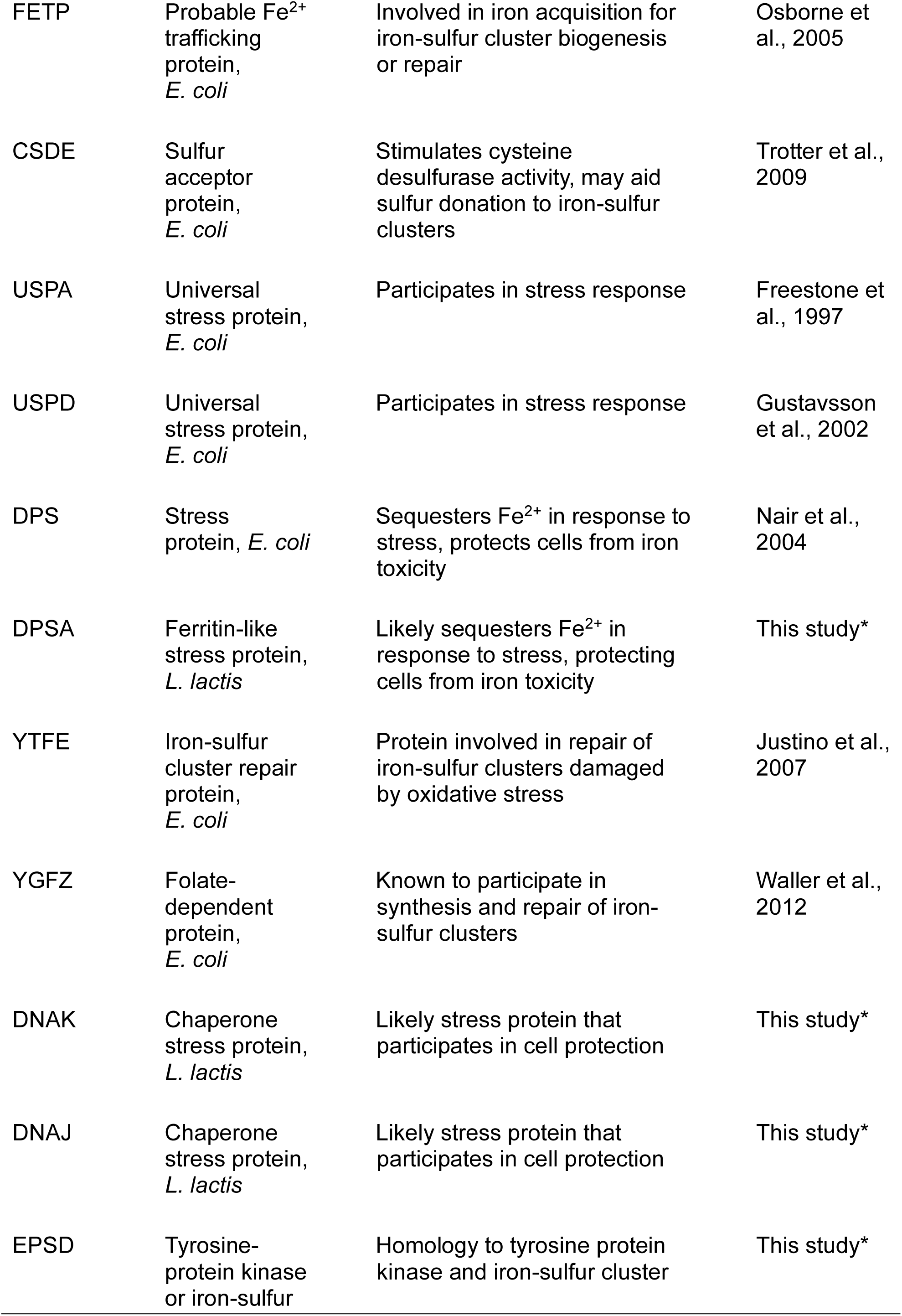
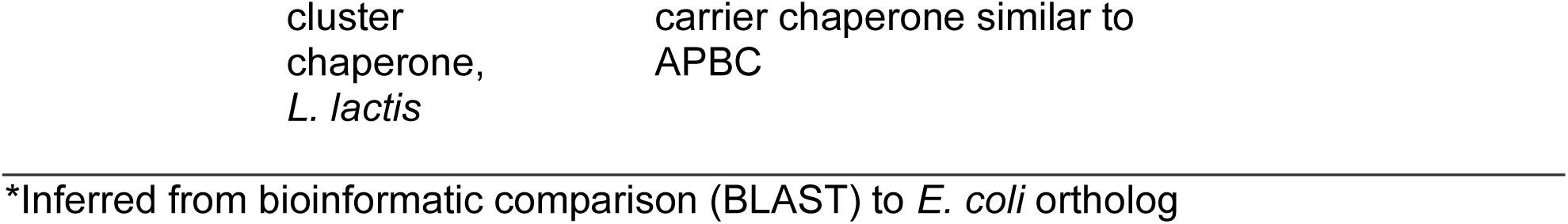
Iron-sulfur cluster accessory genes to co-express

### 2.4 Construction of strains and growth media

*S. cerevisiae* strains in this study were created from CEN.PK2-1C (MATα ura3-52 trp1-289 leu2-3 112 his3Δ1 MAL2-8 SUC2) as described in Supp. Table 1. Deletions, such as *ILV3*, were made using homologous recombination of amplified selection markers. pYZ17 and pYZ84 were amplified by PCR with 75-90 base pair overhangs 5’ and 3’ of the selective marker ORF flanked by loxP sites for marker recycling (with oligonucleotides JDC26-29, Supp. Table 4). *ILV3* was replaced with the drug resistance marker for Nourseothricin (NatMX) and GAL80 with G418 (KanMX). JDCy125 (*ILV3*Δ), JDCy219 (*GAL80*Δ), and other deletion plasmids described in Supp. Table 2 were created by transforming the amplified products and selecting on plates with corresponding drugs. Gene deletions were genotyped via PCR with Thermo Plant Mix PCR to indicate marker presence and removal of gene target ORF. A pooled library was created by first transforming the *ILV3*Δ strain with different BGC CEN plasmids, kept by a TRP marker, via lithium acetate chemical transformation (Gietz et al., 2007). Each strain with the different BGC CEN plasmids were transformed, via electroporation (as previously done; Loock et al., 2023), with the pooled library of DHAD orthologs paired with iron-sulfur cluster accessory genes. Strains used for fermentation were transformed with CEN plasmids containing URA markers and single copy candidate genes. Strains used for fermentations were chassis BGC strains transformed with URA CEN plasmids used to hold single copy versions of the *L. lactis* ilvD (DHAD) and the ferritin-like protein FTNB (from *E. coli*).

Unless specified, yeast were grown on YPD media (20g/L glucose, 10g/L yeast extract, 20g/L peptone and 0.15g/L tryptophan), synthetic complete media (20g/L glucose, 1.5 g/L yeast nitrogen base without amino acids or ammonium sulfate, 5g/L ammonium sulfate, 36mg/L inositol, and 2 g/L amino acid mixture lacking uracil, tryptophan, isoleucine, valine, or leucine), or minimal drop in media (20g/L glucose, 1.5g/L yeast nitrogen base without amino acids or ammonium sulfate, 5g/L ammonium sulfate, 36mg/L inositol, and 1.67mM histidine or leucine).

### 2.5 Complementation assay and nanopore sequencing

Complementation assays took advantage of a valine auxotroph created by *ILV3* deletion. The 4 different yeast libraries (*L. lactis* BGC, *E. coli* BGC, *B. subtilis* BGC, or without BGC) of DHAD orthologs and iron-sulfur cluster accessory genes were pooled by collecting colonies from the selection plates and diluted into liquid Phosphate-Buffered Saline (PBS) solution at 100 OD each. 100 microliters of the pooled libraries were plated onto selective media (synthetic complete media (SC) without uracil, tryptophan, valine, isoleucine, or leucine; minimal drop in media (MM) with histidine or leucine) in triplicate. Colonies were collected and diluted into 1mL of corresponding selective liquid media.

Resuspended yeast cells from colonies were lysed with Zymoprep yeast plasmid miniprep I with zymolyase and isopropanol precipitation. Extracted DNA was amplified via PCR with LongAmp taq polymerase and primers JDC369/370 (Supp. Table 4). Amplified DNA was processed with nanopore sequencing as illustrated in Supp. Fig. 5. Basecalling from raw data and analysis was done as illustrated in Supp. Fig. 6, Python script used for analysis can be found at: https://github.com/jdcortez93/ Avalos_bioinformatics. To calculate log2FC, the representation of the ortholog reads over the total reads in the selective condition were compared to the ortholog reads over total reads in the non-selective condition.

### 2.6 Fermentation and measurement of BCHA production

Fermentations were done at high cell densities similarly to previously described (Zhang et al., 2022). Isolated colonies after transformations were grown overnight at 30°C in 1mL of SC media lacking specific amino acids such as uracil and tryptophan with 2% glucose using sterile 24-well plates. The following day, plates were centrifuged at 3000xg for 3 min, supernatant discarded, and cells resuspended in 1mL of similar SC media with 10% glucose. After incubating for 48 h at 30 °C with 200rpm agitation, plates were again centrifuged at 3000xg for 3 min, and supernatant was saved for analysis. All 24-well fermentation plates were sealed with sterile SealPlate® film to prevent evaporation and cross-contamination between wells. For each genotype, titers represent at least three separate colonies collected from a single transformation.

Isobutanol and isopentanol concentrations were measured by gas-chromatography (GC-MS) using Agilent Infinity instruments. 800 microliters of fermentation supernatant were centrifuged at 17,000xg for 45 min at 4 °C to remove any residual debris. To extract alcohols, supernatant from the centrifuged samples were mixed with hexane at a 1:1 ratio, vortexed for 15min, and centrifuged at 17,000xg for 10 min at 4 °C. Organic phase was analyzed using a DB-WAX UI 0.5 μm gas chromatography column (Agilent, Part No. 122-7033UI, Santa Clara, CA, USA).

### Statistical analysis

Two-way ANOVA analyses were conducted to determine statistical significance of differences observed in isobutanol and isopentanol titers. P-values less than 0.05 are considered sufficient to reject the null hypothesis that the means of two samples are the same.

## 3 Results

### 3.1 Construction of cytosolic BCHA pathway and valine auxotrophy

Previous studies have shown that *S. cerevisiae* can be engineered to produce BCHAs via cytosolic or mitochondrial pathways (Zhang et al., 2022). We focused our analysis on BCHA production with a cytosolic route (Fig. 1A) formed by constitutive expression of prokaryotic acetolactate synthase (ALS, AlsS from *B. subtilis*), modified NADH-dependent ketol-acid reductoisomerase (KARI^P2D1-A1^, ilvC^P2D1-A1^ from *E. coli*), and libraries of dihydroxy-acid dehydratase (DHAD) enzymes and Iron-sulfur cluster related genes from various prokaryotic hosts (Supp. Fig 2B and Table 1). Together, these enzymes synthesize α-ketoisovalerate (α-KIV) in the cytosol, analogous to the mitochondrial pathway involving *ILV2*, *ILV5*, and *ILV3*. Subsequently, α-KIV is converted to isobutanol through the Ehrlich pathway (Park and Hahn, 2019). Because we are testing multiple DHAD orthologs, it is important to know which would be fully functional in a fermentation.

**Figure 2.**
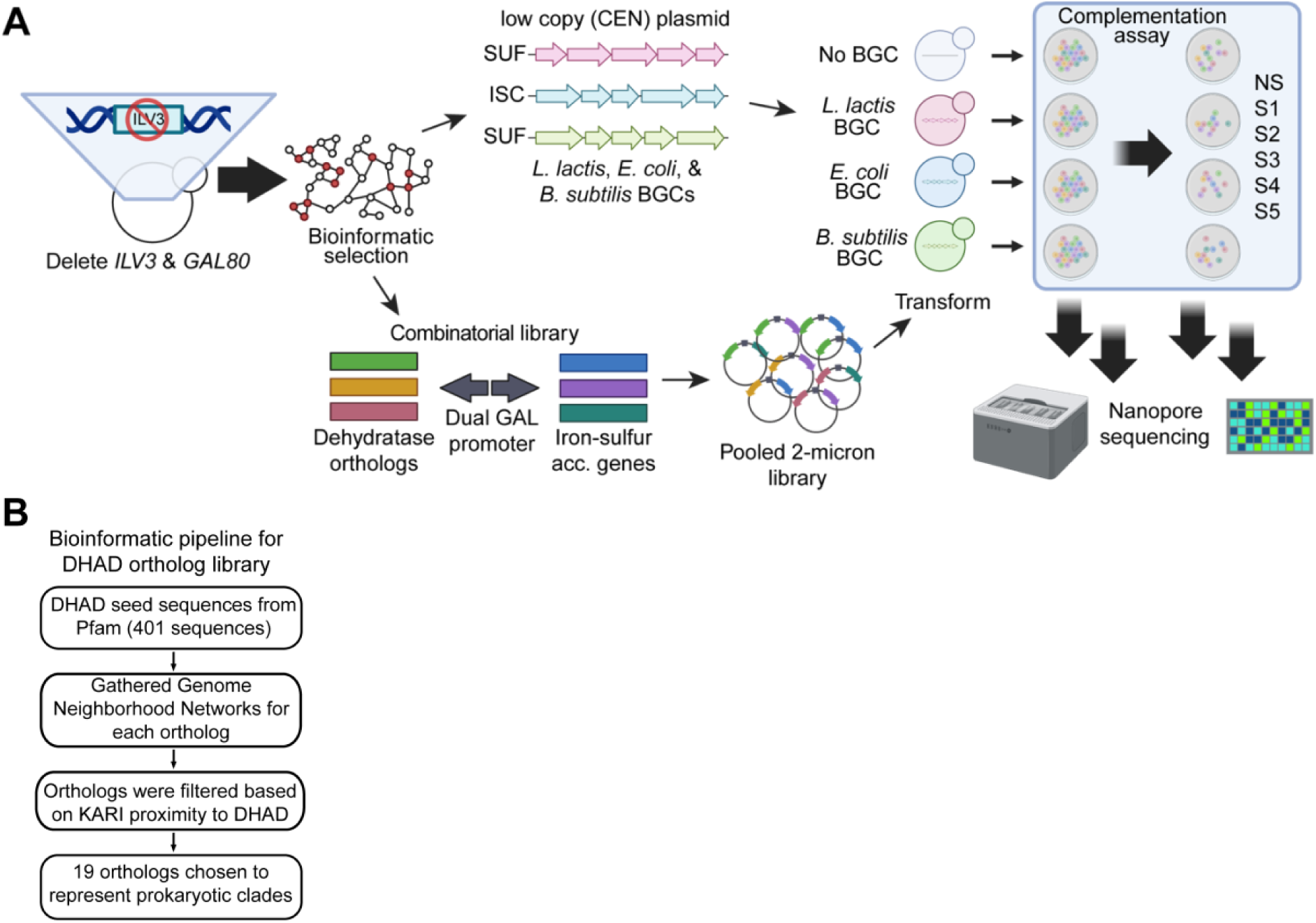
Workflow to test heterologous DHAD activity and generate DHAD ortholog library: (A) Framework of complementation assay. A bioinformatic pipeline and literature review aided in selection of *L. lactis*, *E. coli*, and *B. subtilis* iron-sulfur cluster biosynthetic gene clusters (BGCs). The BGCs were codon-optimized, synthesized, and assembled on low-copy plasmids (CEN) then transformed into yeast with *ILV3* and *GAL80* deleted. Bioinformatics described previously were used to select DHAD orthologs that were codon-optimized and assembled on high-copy plasmids with select iron-sulfur cluster accessory genes on dual GAL promoters. The high-copy (2u) library was transformed into strains with or without different BGCs. Transformants were grown on different media: NS, non-selective synthetic complete (SC) media with no tryptophan or uracil: S1, SC media with no valine; S2, SC media with no valine and isoleucine; S3, SC media with no valine, isoleucine, and leucine; S4, minimal drop in (MM) media with histidine and leucine; S5, MM media with only histidine added. Colonies grown were collected, lysed, and subject to nanopore sequencing. (B) Bioinformatic pipeline to select the 19 DHAD orthologs to assay.

To test the activity of heterologous DHADs (and associated iron-sulfur cluster related genes) we used a valine complementation assay. To achieve this, we generated a valine auxotrophic strain by deleting the endogenous *ILV3,* as done previously (Brat et al., 2012). To validate whether this assay could detect functional DHADs, we cultured a *ΔILV3* strain (JDCy125, Supp. Table 1) in valine-deficient media with or without expression of ilvD from *L. lactis* (*Zhang et al., 2022*). While the wild-type strain grew on synthetic complete media lacking valine, the Δ*ILV3* strain showed no growth (Fig. 1B). However, growth was restored by strong constitutive expression of *L. lactis* ilvD on a low-copy CEN plasmid, confirming functional dependency on cytosolic DHADs. Notably, strains expressing either wild-type or enhanced mutant ilvD^I433V^ (Zhang et al., 2022) on high-copy (2-micron) plasmids exhibited improved growth relative to low-copy plasmid expression (Fig. 1B). Complementation with *L. lactis* ilvD also restored growth in liquid synthetic complete media lacking valine (Supp. Fig. 1). These results indicate that a valine complementation assay is indeed an effective way to screen for functional heterologous DHAD activity.

With this complementation assay, we devised a workflow to test the activity of heterologous DHADs and iron-sulfur cluster-related genes (Fig. 2A). First, we developed a bioinformatic pipeline to identify a library of DHAD orthologs and iron-sulfur cluster BGCs (section 4.2). We generated three different chassis strains expressing iron-sulfur cluster BGCs from *E. coli*, *B. subtilis*, and *L. lactis* using strong promoters and CEN plasmids (Supp. Table 2). Separately, we prepared a library of 2-micron plasmids to express the select DHAD orthologs alone or co-express the select DHAD orthologs paired with iron-sulfur cluster accessory genes individually (Table 1) using a dual GAL promoter that has strong constitutive expression in a *ΔGAL80* strain. This library of plasmids was used to transform the chassis strains with or without BGC genes. Using the complementation assay described above, we plated the transformed libraries in selective media lacking different combinations of BCAAs. Because ILV3p is also involved in the biosynthesis of isoleucine and leucine (Fig. 1A), this assay also works to complement the isoleucine and leucine auxotrophies of the strain (Brat et al., 2012). We collected the selected colonies from each media and sequenced their 2-micron plasmids with nanopore sequencing.

### 3.2 Bioinformatic pipeline and selection of DHAD orthologs and iron-sulfur cluster related genes

To identify a phylogenetically diverse yet experimentally tractable set of DHAD orthologs, we designed a custom bioinformatic pipeline (Fig. 2B). We created a prokaryotic protein database of 371 members from proteomes representing all known prokaryotic organisms in Uniprot (see Methods). We then assembled an alignment-based phylogenetic tree of DHAD orthologs (Fig. 3A) and calculated patristic distances from the *ILV3* sequence. Kernel density estimations of these species’ patristic distances were used to select dehydratase candidates for complementation screening, aiming to have a wide representation of distances (Supp. Fig. 2A).

**Figure 3:**
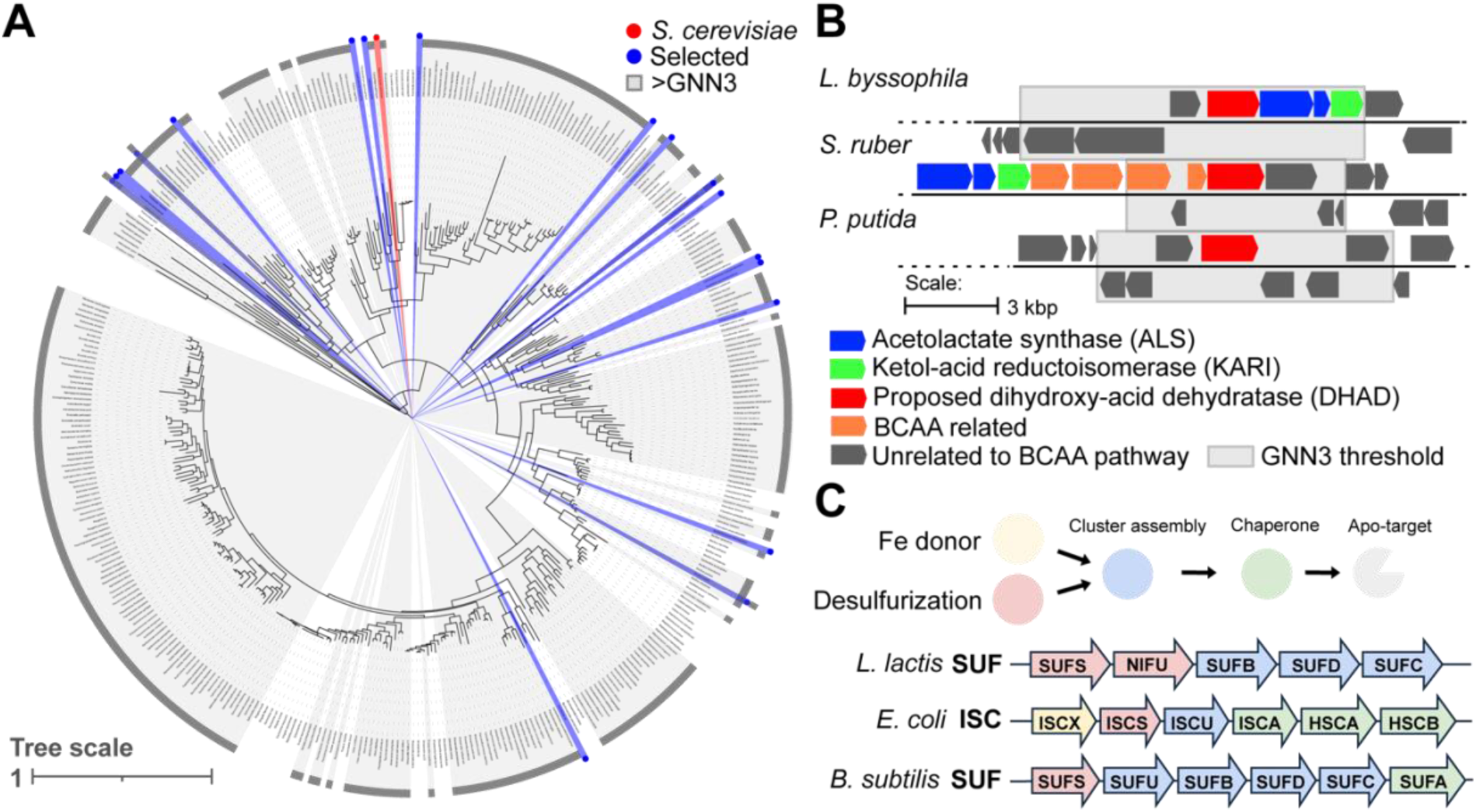
(A) Phylogenetic tree of DHAD orthologs. *S. cerevisiae* is indicated in red; those selected for expression in blue, and in grey (>GNN3) are dehydratases with no KARI in the 3 closest upstream and downstream genomic nearest neighbors. Larger image of tree can be found in Supp. Fig. 5. (B) Bioinformatic analysis of KARI neighbor to DHAD orthologs. Examples of mapping for *L. byssophila, S. ruber*, and *P. putida*. Blue indicates locations of acetolactate synthase (ALS), green is ketol-acid reductoisomerase (KARI), red is the proposed dihydroxy-acid dehydratase (DHAD), orange for BCAA related genes (i.e. leucine biosynthesis), and dark grey are genes not directly related to BCAA biosynthesis. (C) Selection of BGCs based on Fe-S biogenesis operons in *L. lactis*, *E. coli*, and *B. subtilis*.

Our bioinformatic pipeline also distinguishes between dehydratase orthologs that target dihydroxy-acids and those that target other substrates (i.e. phosphogluconate, arabinate, or pentonate; Rahman et al., 2018). By leveraging genomic nearest neighbor (GNN) information of ketol-acid reductoisomerase (KARI), the upstream enzyme in BCAA/BCHA production (Fig. 1A), we cataloged a subset of dehydratases from a list of 371 orthologs in Uniprot (see Methods, Supp. File 1). We hypothesized that close proximity to KARI is indicative of dehydratases likely to target dihydroxy-acid substrates. We included dehydratases that are three genes away from neighboring KARIs (≤GNN3). Based on this value, we propose that 114 dehydratases (31%, Fig. 3A) likely act as DHADs. We reassembled these 114 orthologs in a phylogenetic tree, and trimmed it using Treemmer (Menardo et al., 2018) to 14 orthologs (Fig. 3A, blue) that represent a wide distribution of patristic distances (Supp. Fig. 2) to yeast *ILV3* (Fig. 3A, red). Interestingly, 69% (257 members) of prokaryotes in the database lack a KARI gene within three neighboring genes of a proposed DHAD ortholog (Fig. 3A, >GNN3, grey). While some of these excluded candidates may still be functional DHADs (for example the *Salinibacter ruber* dehydratase, Fig. 3B), applying a threshold of three neighboring genes retained a manageable number of DHAD orthologs for experimental evaluation. As a control, and to expand our diversity, we also included four additional orthologs lacking KARI neighbors (Fig. 3A, blue and grey) and one archaeal dehydratase from *Methanothermobacter thermautotrophicus* (Uniprot ID O27498). These 19 genes were codon-optimized and synthesized for heterologous expression in *S. cerevisiae*.

While the BGCs for *E. coli* and *B. subtilis* have been previously described (Biz and Mahadevan, 2021), the corresponding genes for *L. lactis* have not been reported. Therefore, we also used bioinformatics to identify the likely BGC genes in *L.* lactis based on sequence similarity to the *B. subtilis* SUF family of genes. We did a BLAST search of the BioCyc curated *L. lactis* genome (Fig. 3C and Supp. Fig. 3), excluding transcription factors and genes with transmembrane domains. We synthesized codon-optimized genes from the *E. coli*, *B. subtilis*, and newly proposed *L. lactis* BGC genes (Supp. Table 2). These genes span the four minimal mechanisms for iron-sulfur cluster cofactor biosynthesis and utilization: desulfurization, iron donor, cluster assembly, and chaperone-mediated cluster delivery; belonging to the ISC and SUF gene families (Biz and Mahadevan, 2021).

However, it is challenging to find genes that act as iron donors due to incomplete characterization of genes outside native operons (Biz and Mahadevan, 2021). In our SUF and ISC BGC synthesis, only ISCX from *E. coli* has a proposed iron donor function (Roche et al., 2015). To co-express iron-sulfur cluster accessory genes, some of which may act as iron donors, we identified a list of genes from literature (Table 1), selecting candidates from *L. lactis*, *E. coli*, and *B. subtilis* known to facilitate iron-sulfur cluster function or biogenesis (references in Table 1). These genes represent diverse functions, including ferritin, glutaredoxin, ferredoxin, flavodoxin, universal chaperone or stress response, iron or sulfur trafficking, and general iron-sulfur cluster repair.

To prepare the library we cloned each DHAD ortholog by itself or paired with each iron-sulfur cluster accessory gene with the exception of *L. lactis* DHAD which was left unpaired as a control. Constructs combining DHAD orthologs and iron-sulfur cluster accessory genes were assembled under dual GAL1/GAL10 promoters on high-copy (2-micron) plasmids for strong constitutive expression in Δ*GAL80* strains (Figure 2A).

### 3.3 Selection of functional DHADs and iron-sulfur cluster genes using a complementation assay

BCAA complementation assays revealed multiple candidate DHAD orthologs, BGC genes, and iron-sulfur cluster accessory genes that complement yeast growth in selective media (Supp. Fig. 4). We transformed Δ*ILV3*Δ*GAL80* strains containing the partial cytosolic BCHA pathway (ALS and NADH-dependent KARI^P2D1-A1^) with CEN plasmids containing different BGCs (from *L. lactis*, *E. coli*, or *B. subtilis,* empty plasmid; strains JDCy221-JDCy224) to generate the three chassis strains and a negative control. We then transformed these strains with a combinatorial library of DHAD orthologs and iron-sulfur cluster accessory genes on 2-micron plasmids, obtaining at least 7 x 10^3^ colonies (approximately 10x library coverage) on SC-URA-TRP. We pooled the transformed colonies and replated them on complementation media lacking valine (S1); valine and isoleucine (S2); valine, isoleucine, and leucine (S3); or minimal media drop-in conditions (S4 with histidine and leucine, S5 with only histidine). With the exception of the *L. lactis* BGC, under the most stringent conditions (S3 and S4) every library showed colonies growing in all complementation media (Supp. Fig. 4A). Based on the number of colonies that grew in non-selective media, we estimate that the number of surviving colonies in the different selective media range between <1% - 8%, revealing that libraries with the *B. subtilis* BGC survived best in most conditions (Supp. Fig. 4B).

**Figure 4.**
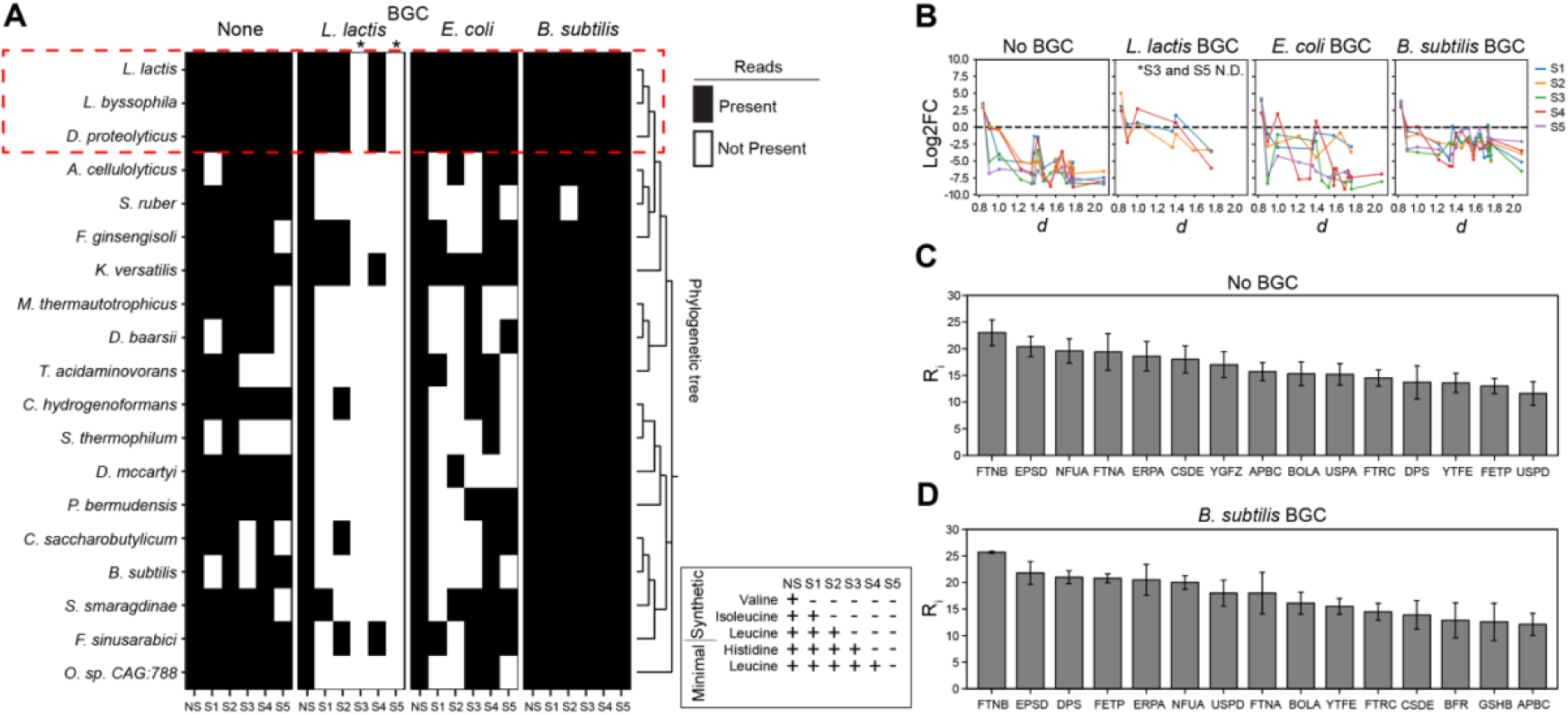
**Read results from complementation assays**: (A) Nanopore reads of candidate DHAD orthologs. NS, non-selective media; S1, synthetic complete media (SC) -valine + 2% glucose; S2, SC -valine, isoleucine + 2% glucose; S3, SC -valine, isoleucine, leucine; S4, minimal drop in media (MM) +histidine, leucine; S5, MM +histidine. (B) Log2FC of DHAD ortholog (represented by dots) read change between NS and selective media (S1-S5; blue, orange, green, red, and purple lines) indicated by *d,* patristic distance from yeast *ILV3*. Values with log2FC of 0 were not plotted. (C) Assisting iron-sulfur cluster accessory genes with no BGC and (D) *B. subtilis* BGC. Ranking score of the most iron-sulfur cluster accessory gene reads present with DHAD orthologs less than 1 patristic distance. FTNB, the ferritin-like gene from *E. coli*, is most present in reads. R_i_ indicates ranking score; highest values are most frequent in the sample. *d* indicates the patristic distance of the ortholog from yeast *ILV3*. * Indicates media that yielded no colonies.

**Figure 5:**
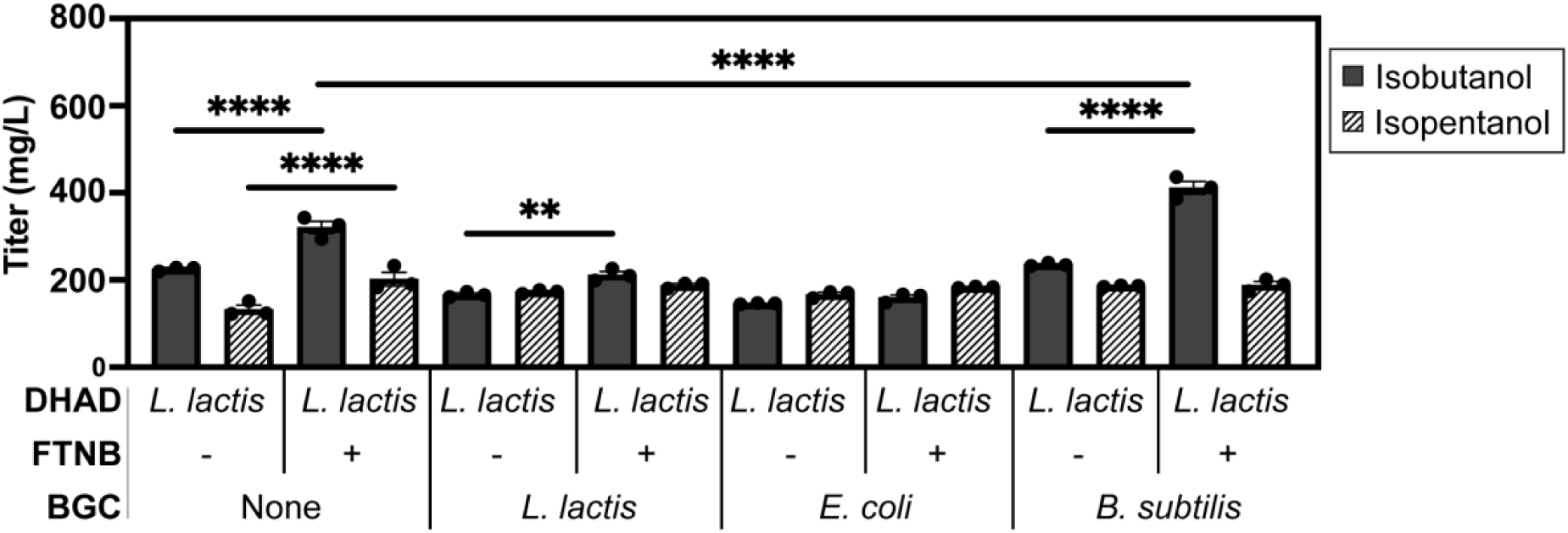
Co-expression of L. lactis DHAD with iron-sulfur cluster related genes. Fermentation with best DHAD ortholog, BGC, and iron-sulfur cluster accessory gene. FTNB, the ferritin-like gene from *E. coli*, is able to improve isobutanol and isopentanol titer without a BGC present. However, the *B. subtilis* BGC, in combination with FTNB made the most significant impact on titer of isobutanol. **P < 0.01 ****P < 0.0001.

Next, we sequenced the active DHAD orthologs and accompanying iron-sulfur cluster accessory genes found in colonies that grew on the selective media (Supp. Fig.

4 and 6). Direct nanopore sequencing of the pooled DNA library prior to transformation confirmed the presence of every synthesized gene (Supp. Fig. 7). For transformed cells that survived selection, we performed PCR amplification then nanopore sequencing using barcoded primers flanking the dual GAL promoter cassette (see Methods for details). In all libraries, greater than 20 reads for each DHAD ortholog were detected in non-selective media (Supp. Fig. 7A). DHAD orthologs with patristic distances less than 1.0 from yeast *ILV3* (*L. lactis*, *L. byssophila*, and *D. proteolyticus*) were consistently detected in all selective conditions even without co-expressing BGCs (Fig. 4A, red segmented box), indicating their likely functional DHAD activity (Supp. Fig. 7 and 8). Interestingly, the *K. versatilis* ortholog, an outlier with patristic distance higher than 1.4, was also present in all selective media (except S3 for cells containing the *E. coli* BCG) and is likely an active DHAD (Fig. 3A). Nonetheless, DHAD orthologs closest in patristic distance to yeast *ILV3* are probably functional enough to provide growth and possible BCHA production.

Strains expressing the *B. subtilis* BGC had the most universal impact on DHAD ortholog activity (Fig. 4A: *B. subtilis* BGC). In the presence of *B. subtilis* BGC, every ortholog tested yielded at least one sequencing read under most selective conditions. Orthologs from *A. cellulolyticus*, *F. ginsengisoli*, *M. thermautotrophicus*, *D. baarsii*, *C. hydrogenoformans*, *P. bermudensis*, *B. subtilis*, and *F. sinusarabici* had sequencing reads in every selective media tested. This suggests that these orthologs are also functional DHADs in yeast cytosol in the presence of the *B. subtilis* BGC.

To depict enrichment, we calculated the log2 fold change (log2FC) of DHAD orthologs (Methods) ordered by patristic distance (Fig. 4B) in selective media (S1-S5) compared to non-selective media (NS). With or without BGCs, *L. lactis* (distance 0.83), *L. byssophila* (distance 0.90), and *D. proteolyticus* (distance 1.00) orthologs had the highest log2FC values with many being positive (>0). However, the *L. lactis* ortholog, with and without the *B. subtilis* BGC, was most robust with the highest positive log2FC values (>2.5) across all selective media types (S1-S5). Analysis of iron-sulfur accessory gene reads from colonies also containing the most successful DHAD orthologs (patristic distance ≤1) revealed associated iron-sulfur cluster accessory genes that aid cytosolic DHAD function (Fig. 4C, D).

The co-expressed iron-sulfur cluster accessory genes likely facilitate functionality of many active DHAD orthologs. Several of these genes were found in the sequencing reads derived from strains that survived selection conditions, which we ranked based on their read frequencies across selective media (Supp. Fig. 10). When analyzing the DHADs most closely related to *ILV3* (Fig. 4A, dashed red box), we found that FTNB (an *E. coli*-derived ferritin-like gene) was the most frequently co-expressed accessory gene in strains with *B. subtilis* BGC or without (Fig. 4C, D). Suggesting that expression of *B. subtilis* BGC and *E. coli* FTNB genes likely enhance the activity of a co-expressed *L. lactis* DHAD and therefore BCHA production.

### 4.4 BCHA production by overexpressing iron-sulfur cluster-related genes

Based on our previous results we hypothesized that co-expression of the *B. subtilis* BGC, FTNB and ilvD from *L. lactis* (the most active DHAD we found) with a complete BCHA biosynthetic pathway (Fig. 1A) would enhance the production of isobutanol and isopentanol. To test this, we transformed Δ*ILV3* chassis BGC strains (i.e. *B. subtilis* BGC) containing a partial cytosolic BCHA pathway (Bs_AlsS and Ec_ilvC^P2D1-A1^) with *URA CEN* plasmids containing *L. lactis* ilvD with or without FTNB. The parent strains also overexpress *AFT1*, a gene involved in iron homeostasis previously shown to improve BCHA production (Zhang et al., 2022). We found that FTNB significantly improves isobutanol and isopentanol production in most strains. In the absence of an iron-sulfur cluster BGC, co-expression of FTNB with the *L. lactis* DHAD ortholog on a CEN plasmid resulted in an average isobutanol titer of 321mg/L, a 1.4-fold increase compared to expression of *L. lactis* DHAD alone. FTNB co-expression also gave an average isopentanol titer of 202mg/L, a 1.5-fold increase to *L. lactis* DHAD alone. Interestingly, individual expression of the *L. lactis*, *E. coli*, and *B. subtilis* BGC alone did not significantly improve isobutanol or isopentanol production when using the DHAD from *L. lactis*. However, co-expression of FTNB with the *L. lactis* BGC significantly increased isobutanol production by 1.2-fold compared to co-expressing the *L. lactis* BGC alone. The highest titer observed (achieved when co-expressing of the *B. subtilis* BGC with FTNB) was an average isobutanol titer of 412mg/L, 1.8-fold greater than expression without any BGC or iron-sulfur cluster accessory gene. This significantly enhanced production indicates a likely synergistic interaction between the *B. subtilis* BGC and FTNB activities.

## 5 Discussion

In this work, we analyzed how a set of heterologous iron-sulfur cluster related genes contribute to the enzymatic function of cytosolic dihydroxy-acid dehydratase (DHAD) orthologs in *S. cerevisiae*. We utilized a BCAA complementation strategy to screen a library of iron-sulfur cluster biosynthetic gene clusters (BGCs) and accessory genes (Fig. 2A). We found that DHAD orthologs with the closest phylogenetic relationship (patristic distance) to yeast *ILV3* had the greatest number of recovered sequences and most robust growth through complementation activity (red box in Fig. 4A), with the ortholog from *L. lactis* having the closest relationship and showing the strongest sequence enrichment. We also found that strong constitutive expression of the *B. subtilis* BGC improved activity for most DHAD orthologs tested, particularly when co-expressed with the ferritin-like iron-binding gene FTNB from *E. coli*.

A common limitation in several biosynthetic pathways for chemical production in yeast involving heterologous iron-sulfur cluster dependent enzymes is the appropriate availability and delivery of these cofactors for proper enzymatic function (Biz and Mahadevan, 2021). For example, in addition to the DHADs for isobutanol production examined in this study, the iron-sulfur cluster dependent enzymes D-xylonate dehydratase (XylD) in xylose catabolism, 2C-methyl-D-erythritol-2,4-cyclodiphosphate (IspG) or 4-hydroxy-3-methylbut-2enyl diphosphate reductase (IspH) in the methylerythritol phosphate (MEP) pathway, all face limited function in yeast, probably due to low iron-sulfur cluster availability and delivery (Carlsen et al., 2013; Partow et al., 2012; Wasserstrom et al., 2018; Salusjärvi et al., 2019). This is consistent with studies suggesting that native machineries are inefficient at recognizing heterologous apo-enzymes (Carlsen et al., 2013; Partow et al., 2012; Kirby et al., 2016). Yeast and many eukaryotes have dedicated iron-sulfur cluster assembly machineries that involve sequestering iron and sulfur, assembling them into clusters, and delivering them to the correct apo-enzyme (Biz and Mahadevan, 2021). Furthermore, these mechanisms are divided between the mitochondria and cytosol, requiring the export of iron-sulfur cluster intermediates from mitochondria to the cytosol (Pandey et al., 2019; Lill and Freibert, 2020). In the mitochondria-lacking prokaryotes, iron-sulfur cluster BGCs such as ISC and SUF are not compartmentalized and would give a more efficient process for yeast cytosolic enzyme activity (Garcia et al., 2022; Biz and Mahadevan, 2021).; Biz and Mahadevan, 2021).

Poor availability and transport of iron-sulfur clusters to the cytosolic apo-DHAD likely have been constraining enzyme activity and making the cytosolic BCHA pathway less successful (Gambacorta et al., 2022). Indeed, our results confirm that addition of the heterologous *B. subtilis* SUF BGC universally improves activity for all DHAD orthologs examined (Fig. 4A), probably due to enhanced cytosolic iron-sulfur cluster assembly. In the SUF pathway, sulfur would be provided by SUFS’s desulfurization activity, the SUFBDC complex assembles the iron-sulfur cluster, and SUFA/ERPA deliver the cluster to the apo-proteins. However, the synthesized *B. subtilis* SUF BGC lacks a clear iron-delivery component, as no gene found in the operon is predicted to do so. Consequently, we included a variety of potential iron donors in our iron-sulfur cluster accessory gene library. The ferritin-like gene FTNB emerged as the most successful candidate to co-express. When expressed in yeast, FTNB, known to maintain bacterial iron homeostasis during oxidative stress (Pourciau et al., 2019), may sequester cytosolic iron and shuttle it to the *B. subtilis* SUFBDC complex. This proposed iron-delivery role would explain why significant increases in BCHA production occurred only when the *B. subtilis* BGC was co-expressed with FTNB (Fig. 5). Additionally, FTNB might participate in the repair of damaged Iron-sulfur clusters (Troxell and Hassan, 2013), potentially explaining why FTNB alone significantly enhanced BCHA titers even in the absence of an iron-sulfur cluster BGC (Fig. 5).

As shown in Fig. 4A, predicted BGCs from *L. lactis* did not substantially enhance DHAD activity as we hypothesized. We found instead that the *B. subtilis* SUF BGC was most beneficial to *L. lactis* and most DHAD ortholog activities. The observation that the *L. lactis* BGC was not most beneficial to the DHAD from the same species contrasts with previous findings in *E. coli* where the probability of functional expression of heterologous Fe-S enzymes and the Fe-S proteins networks to maintain them was correlated with the phylogenetic distance between them (D’Angelo et al., 2022). One explanation for our differing results may be the potential toxicity from strong constitutive expression of the entire *L. lactis* BGC, which inhibited colony formation under stringent selective conditions (Supp. Fig 4, S3 and S5). Alternatively, the *L. lactis* BGC may have lower functionality since it was bioinformatically predicted due to limited available characterization of iron-sulfur cluster pathways in this organism.

Additional genes identified from our complementation screen remain promising candidates to co-express and improve DHAD activity in BCHA fermentations. Among the top-ranked accessory genes identified with B. subtilis BGC or without are EPSD, FTNA, FETP, DPS, NFUA, and ERPA (Fig. 4C,D). EPSD, a gene from *L. lactis*, is involved in tyrosine-protein kinase and could potentially phosphorylate iron-sulfur cluster-associated proteins (Elsholz et al., 2014). FTNA (*E. coli*), FETP (*E. coli*), and DPS (*L. lactis*) interact with iron and likely promote iron sequestration to biogenesis complexes (Stillman et al., 2001; Osborne et al., 2005; Nair et al., 2004). NFUA and ERPA (*E. coli*) are proposed iron-sulfur cluster chaperones and could improve iron-sulfur cluster delivery to apo-DHADs. We hypothesize that co-expressing these genes, with increased expression of the *L. lactis* DHAD ortholog, would further improve iron availability, iron-sulfur cluster delivery, and overall cytosolic DHAD performance to give an increased production of BCHA products such as isobutanol and isopentanol.

## Declaration of interest

### Declaration of Generative AI and AI-assisted technologies in the writing process

During the preparation of this work the authors used chatGPT4.5 in order to improve readability and language of the work. After using this tool/service, the authors reviewed and edited the content as needed and take full responsibility for the content of the publication.

## Author contributions

JDC, conceptualization, data curation, formal analysis, investigation, methodology, software/coding, writing – original draft, review & editing; JLA, investigation, methodology, project administration, supervision, writing – original draft, review & editing.

## Acknowledgements

The authors would like to acknowledge Jef Boeke and Aleksandra Wudzinska for use of their nanopore facility and valuable feedback. We also thank Yasuo Yoshikuni and the entire team at the JGI DNA synthesis program for their valuable insight in making the DNA library for this study. We also acknowledge the contribution to ideas from Mina Takegami and Joyce Mo. Finally, we thank all the members of the Avalos lab for their assistance in editing and brainstorming ideas.

## Supplemental Tables

**Supplementary Table 1.**
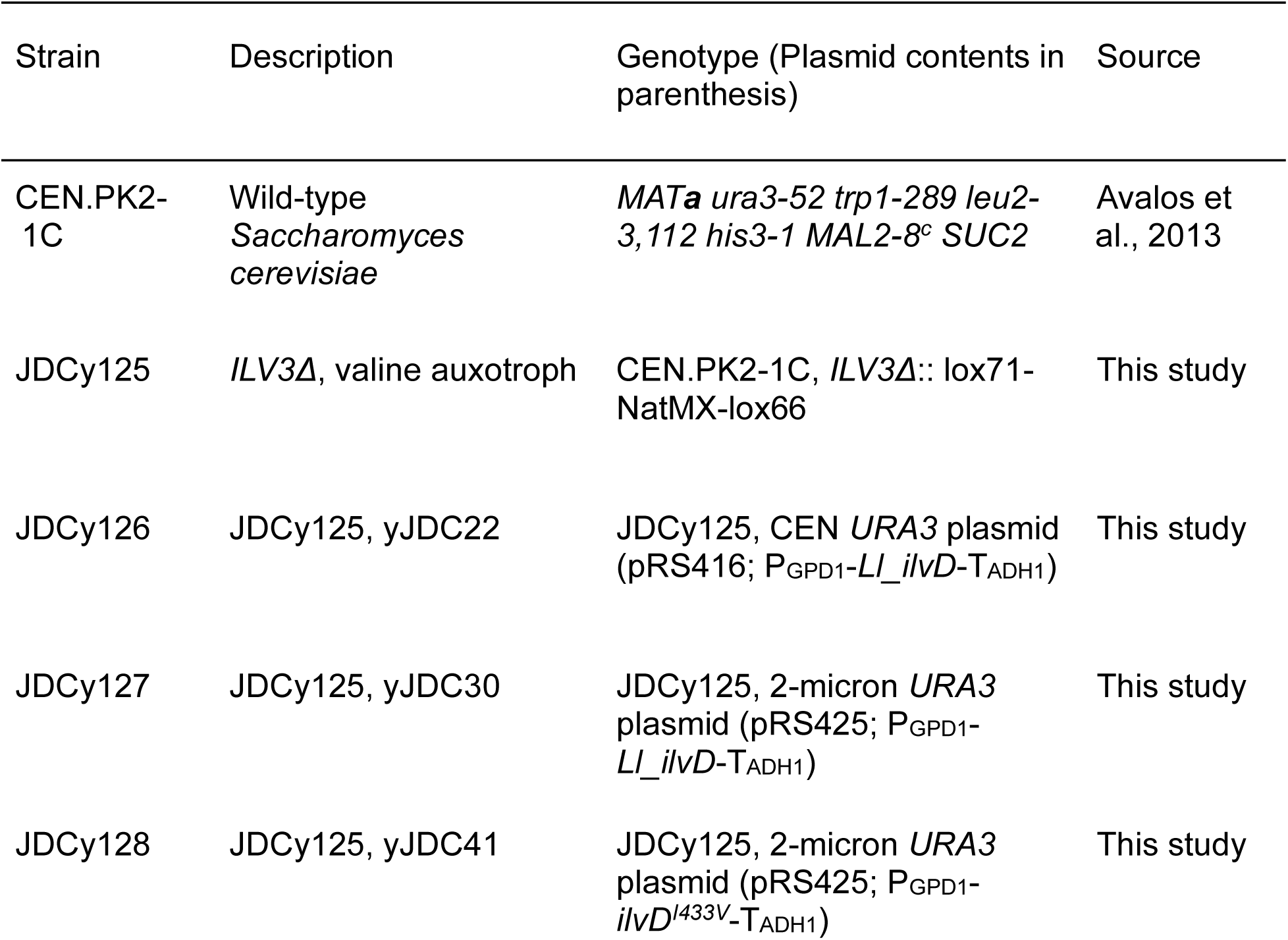
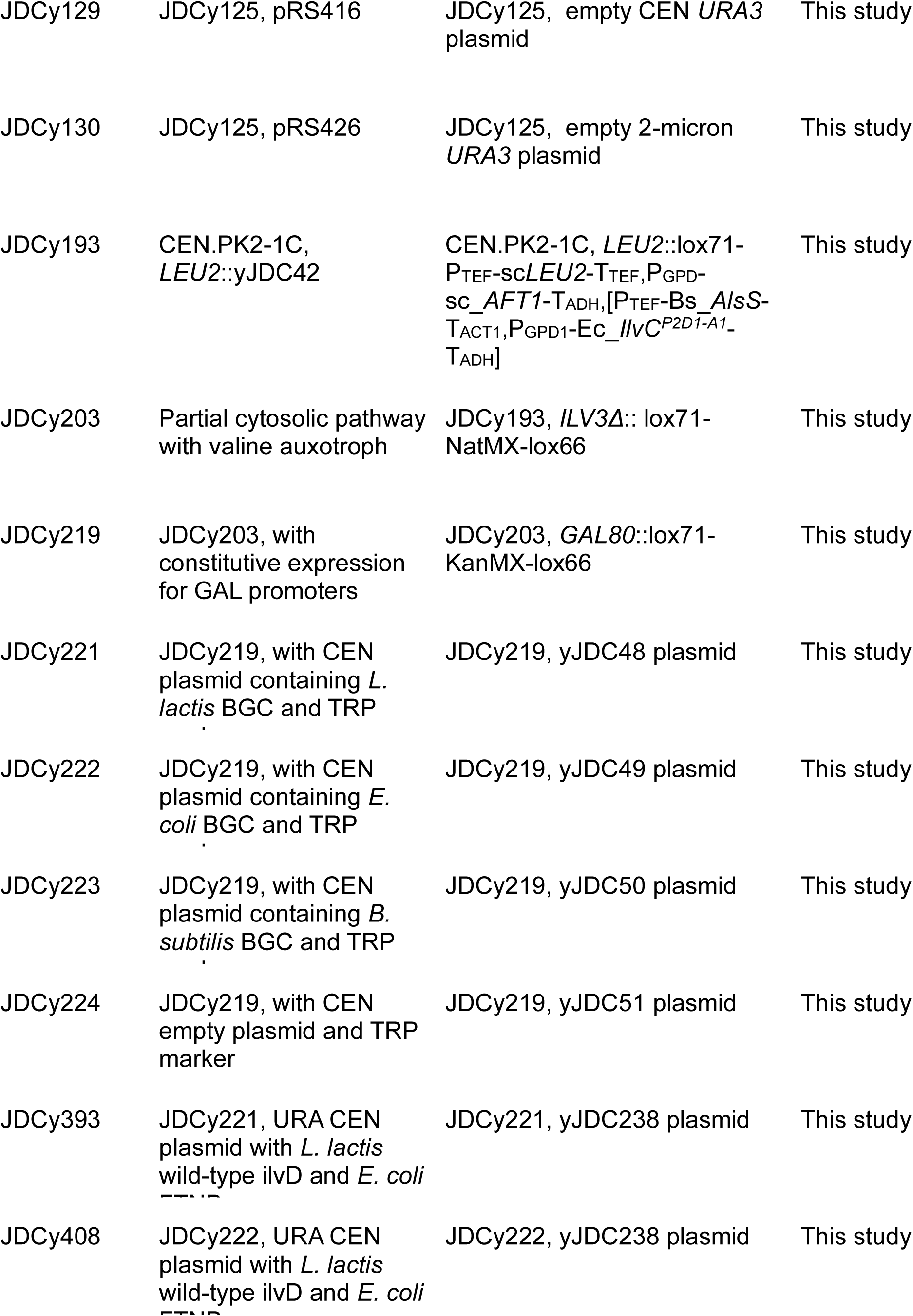
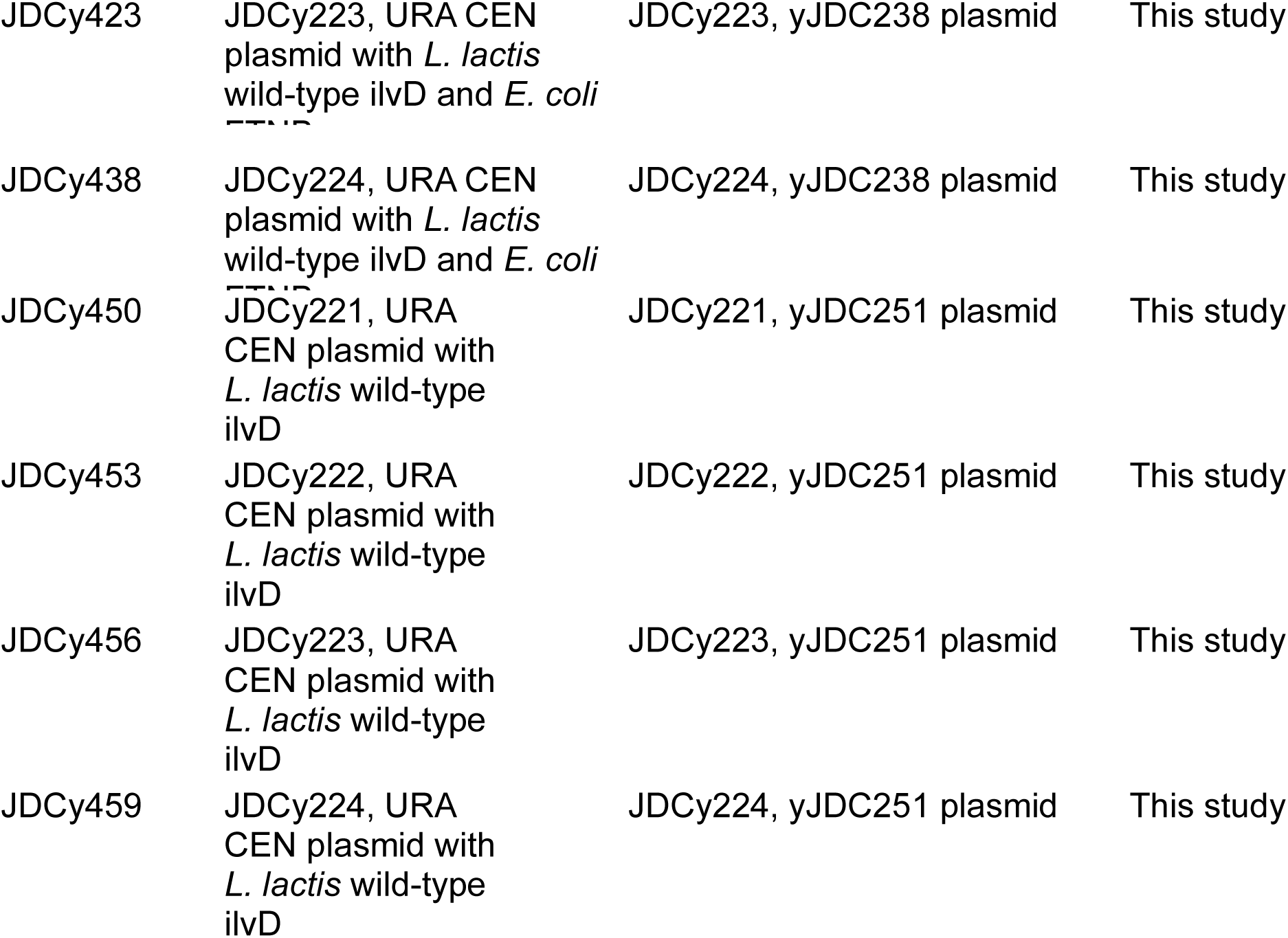
Yeast strains used in this study.

**Supplementary Table 2.**
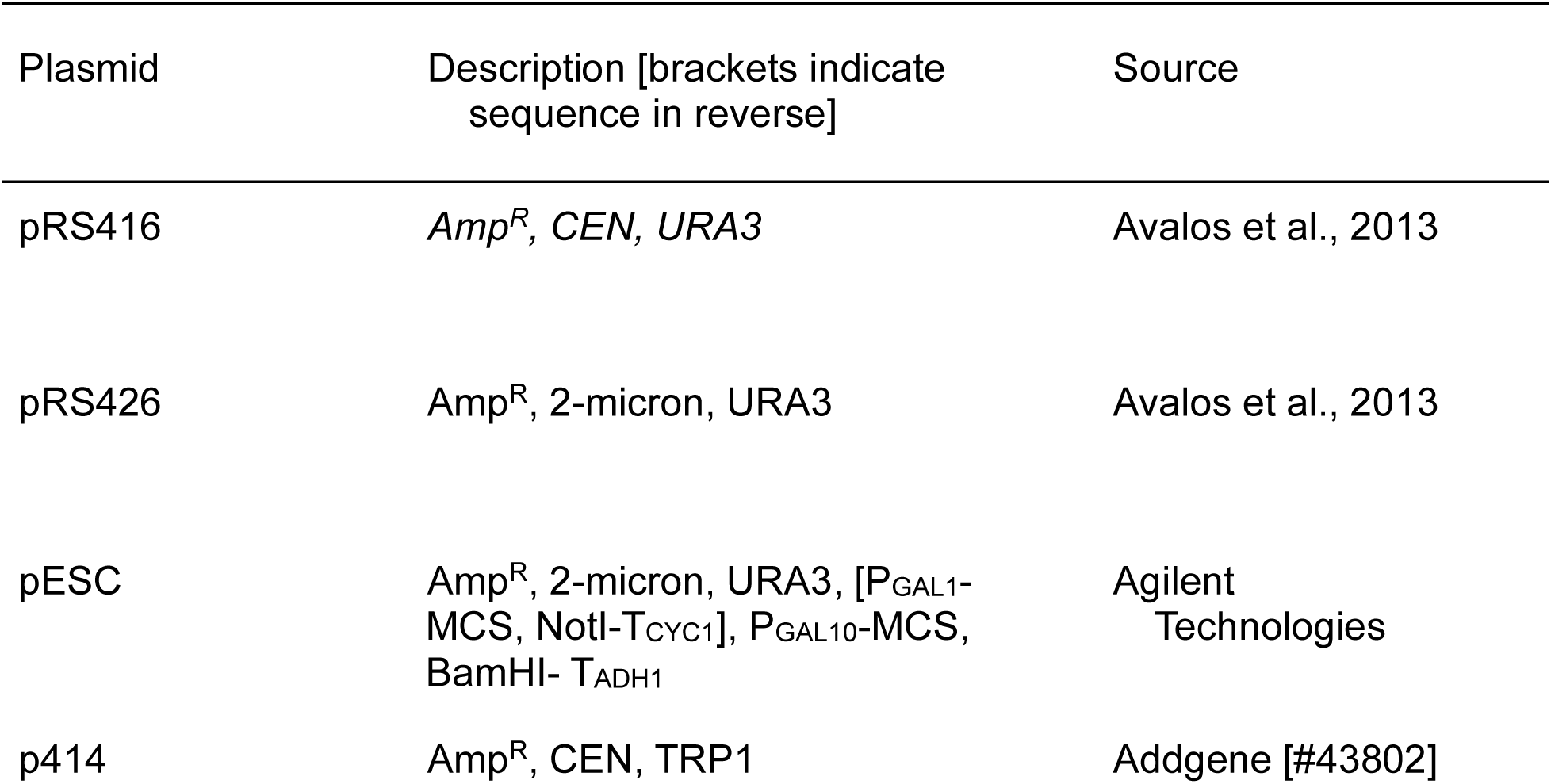
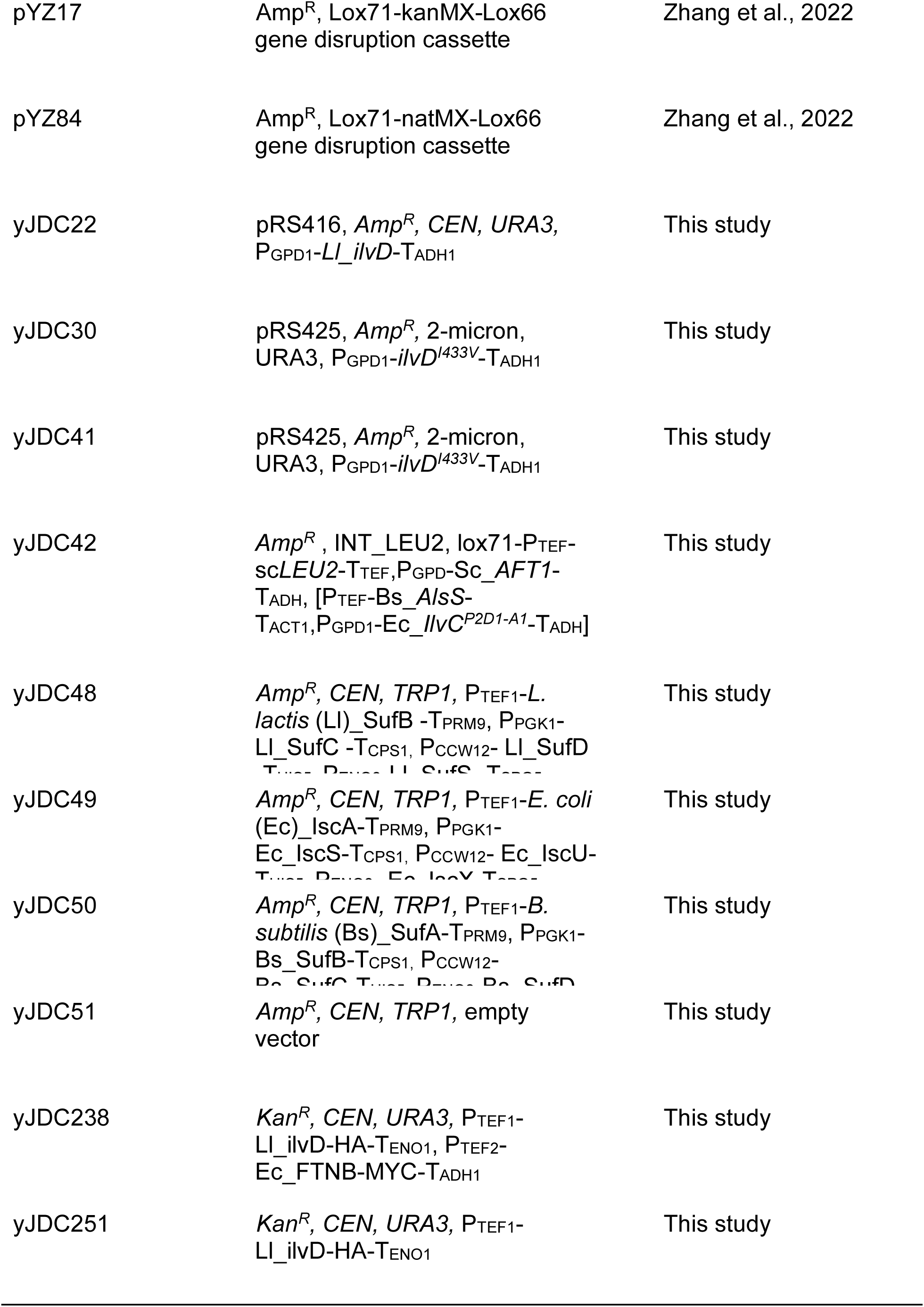
Plasmids used in this study.

**Supplementary Table 3:** E-values of L. lactis BGC search.

| *L. lactis*<br>gene name | *B. subtilis*<br>gene name | Proposed function (in *B. subtilis*) | E-value |
| --- | --- | --- | --- |
| YseI | SUFS | Cysteine desulfurase, acts as initial step in<br>SUF-like Fe-S cluster assembly | 8e-135 |
| NifU | SUFU | Zinc-dependent sulfurtransferase. Transfers<br>sulfur from SufS to SufB. | 1.9e-37 |
| YseF | SUFB | Part of the SufBCD complex. Contributes to<br>assembly or repair of Fe-S clusters. | 2e-111 |
| YsfB | SUFC | Part of the SufBCD complex. Contributes to<br>assembly or repair of Fe-S clusters. | 6e-91 |
| YsfA | SUFD | Part of the SufBCD complex. Contributes to<br>assembly or repair of Fe-S clusters. | 9e-14 |
| YkiE | YqkB | FeS cluster biogenesis domain-containing<br>protein. Possible Fe-S cluster carrier. | 0.0011 |
| YjaF | SUFT | Fe-S protein maturation auxiliary factor.<br>May function as a Fe-S cluster carrier. | 7.3e-41 |

**Supplementary Table 4.**
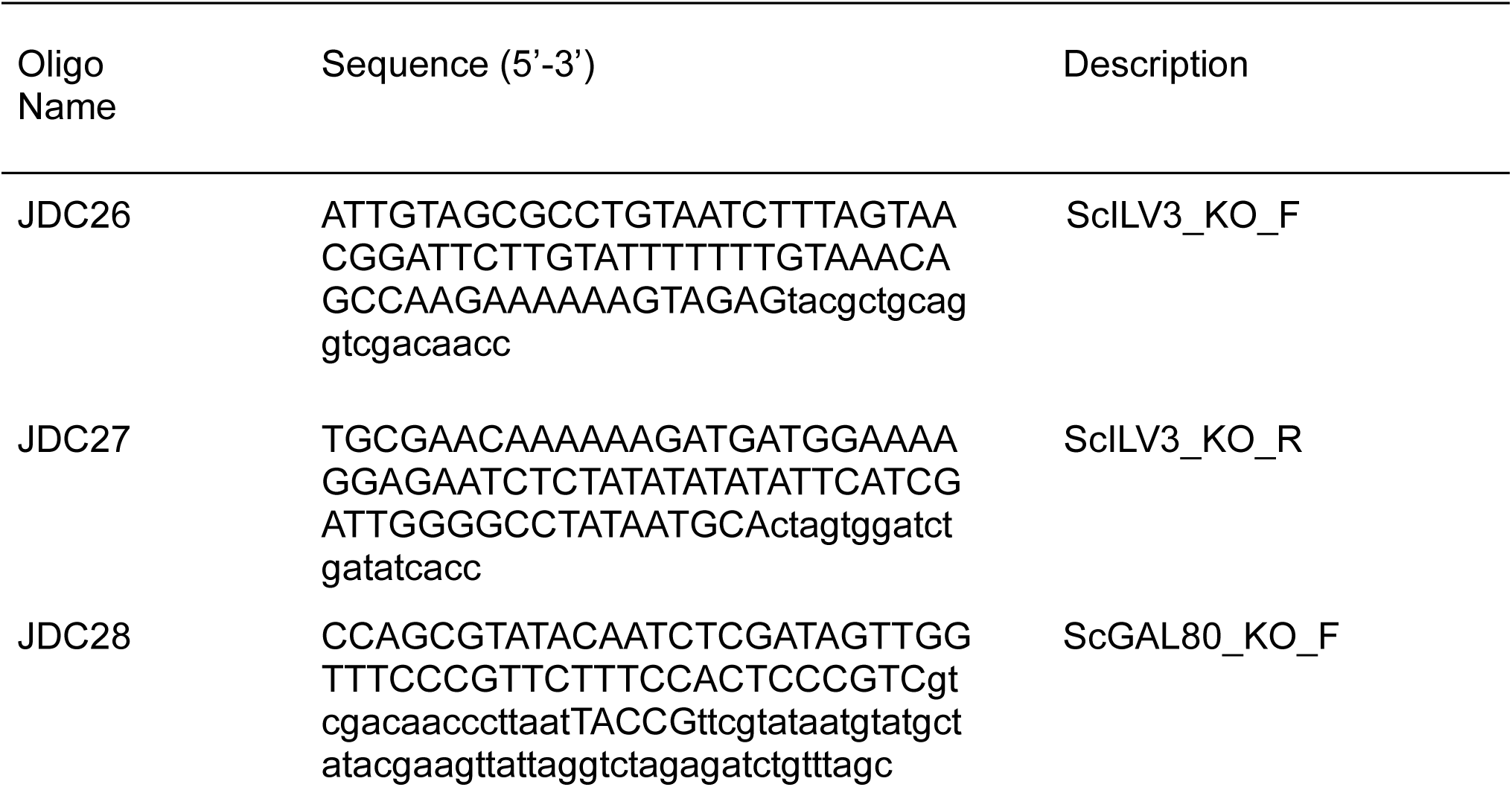
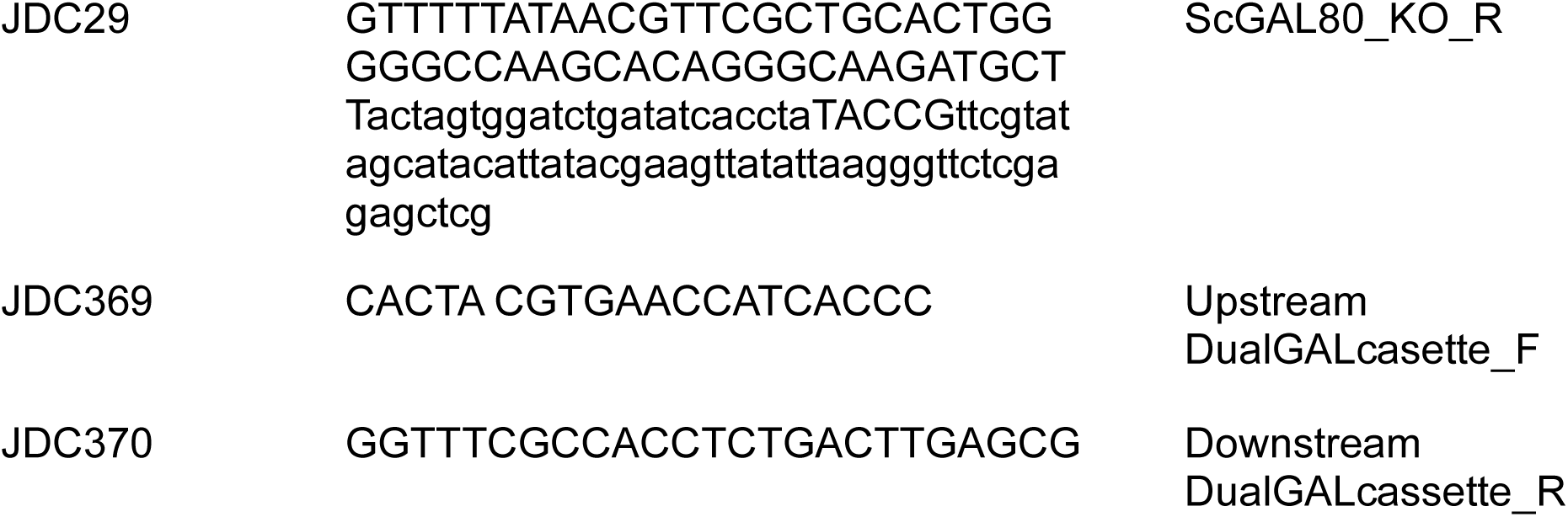
Oligonucleotides used in this study.

## Supplemental Figures

**Supplementary figure 1:**
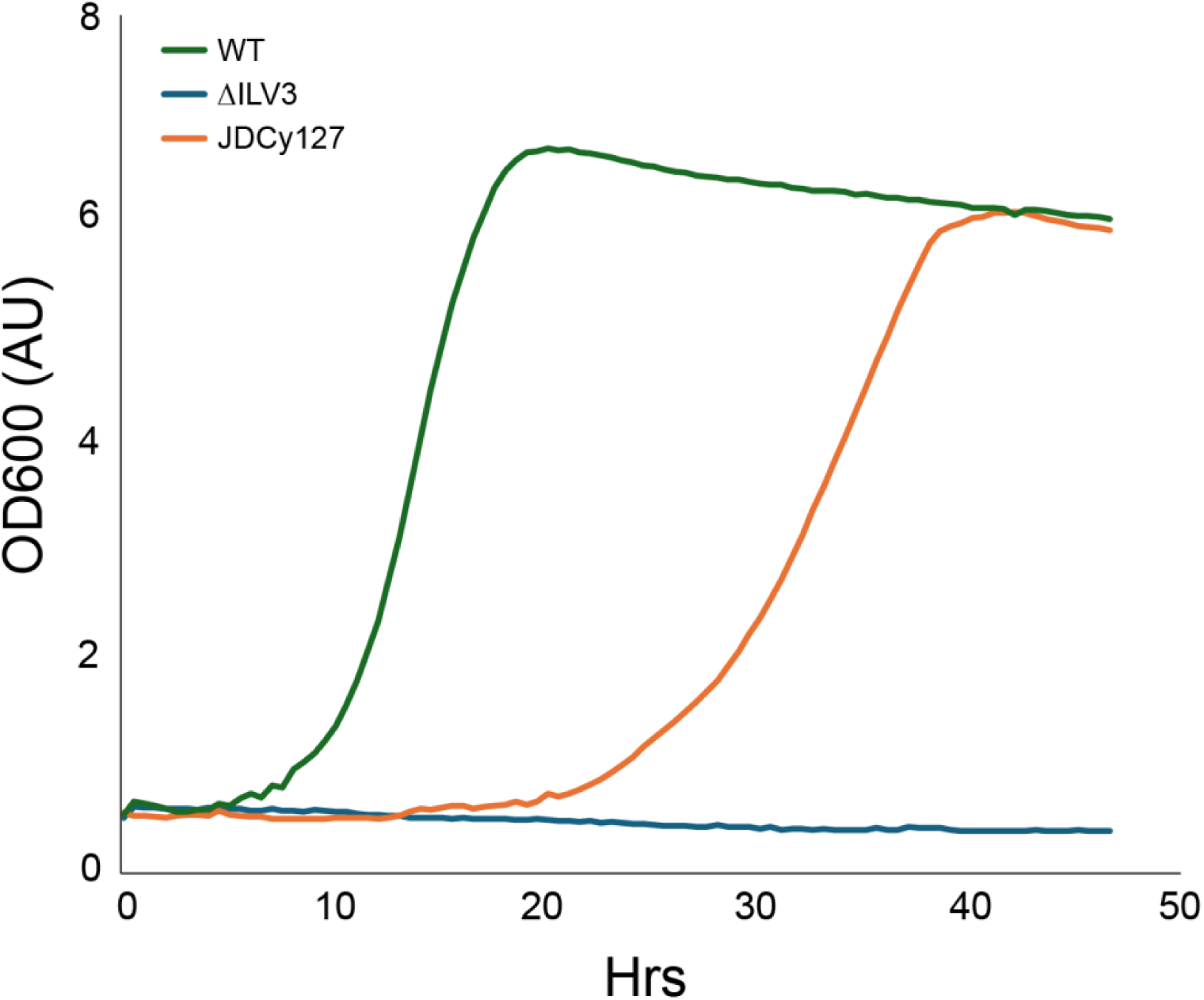
*L. lactis* DHAD rescues Δ*ILV3* in synthetic complete (SC) without valine and uracil. Growth curve in liquid media SC - valine, uracil +2% glucose, heterologous expression of *L. lactis* DHAD is able to rescue growth. WT (green) represents the wild-type CEN.PK. strain without an *ILV3* deletion, *ΔILV3 (*blue) represents CEN.PK. with *ILV3* deleted and empty plasmid, JDCy127 (orange) represents *ΔILV3* with high copy plasmid of strong constitutive expression (pGPD1) for ilvD from *L. lactis*.

**Supplementary figure 2:**
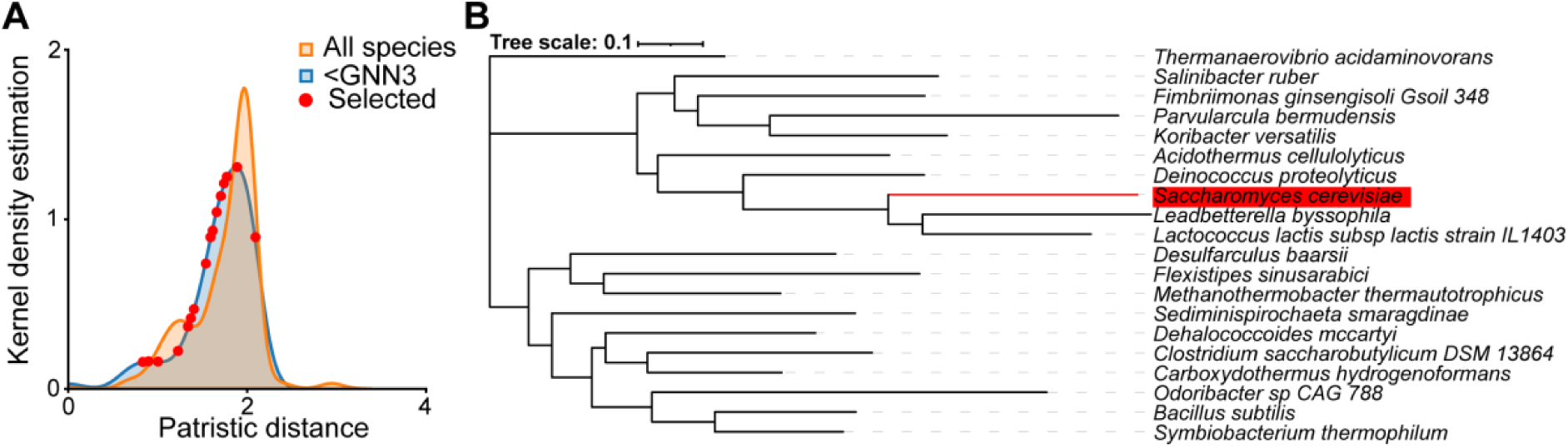
(A) Patristic distance distribution. Distribution of patristic distance from *S. cerevisiae ILV3*, represented in orange, indicates a majority from 0 to 2. Species, represented in blue, with at least 1 KARI in the 3 upstream and downstream genes from the proposed DHAD maintain similar distribution but shifted slightly closer in patristic distance. Red dots represent the dehydratase orthologs selected to assay. (B) Phylogenetic tree of DHAD ortholog candidates. Representative phylogenetic tree of select dehydratases in relation to *S. cerevisiae ILV3* (highlighted in red). Closest relatives are dehydratases from *L. byssophila* and *L. lactis*.

**Supplementary figure 3:**
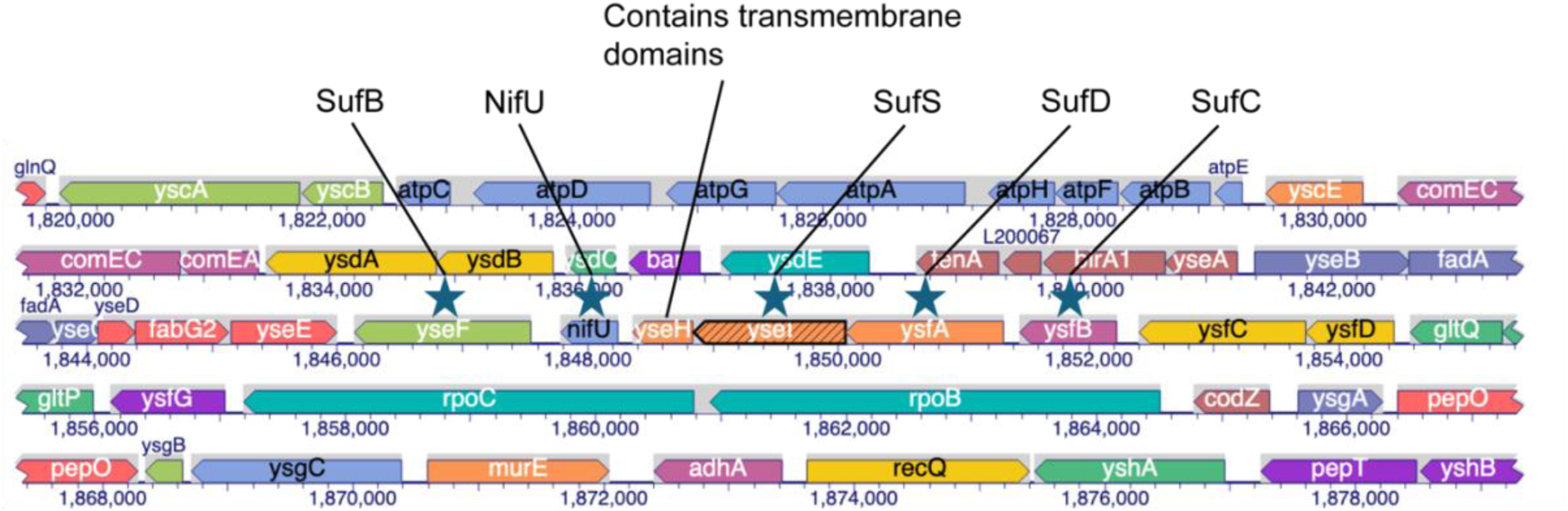
*Lactococcus lactis* genome with iron-sulfur cluster BGC orthologs. Gene mapping created from BioCyc genomic data of *L. lactis*. BGC genes from the *SUF* pathway were BLAST searched. Majority of significant hits found were within the window shown, all with e value lower than 10^-10^. SufB matched with yseF (evalue = 2e-111), sufC to ysfB (2e-91), sufD to ysfA (9e-14), sufS to yseI (8e-135).

**Supplementary figure 4:**
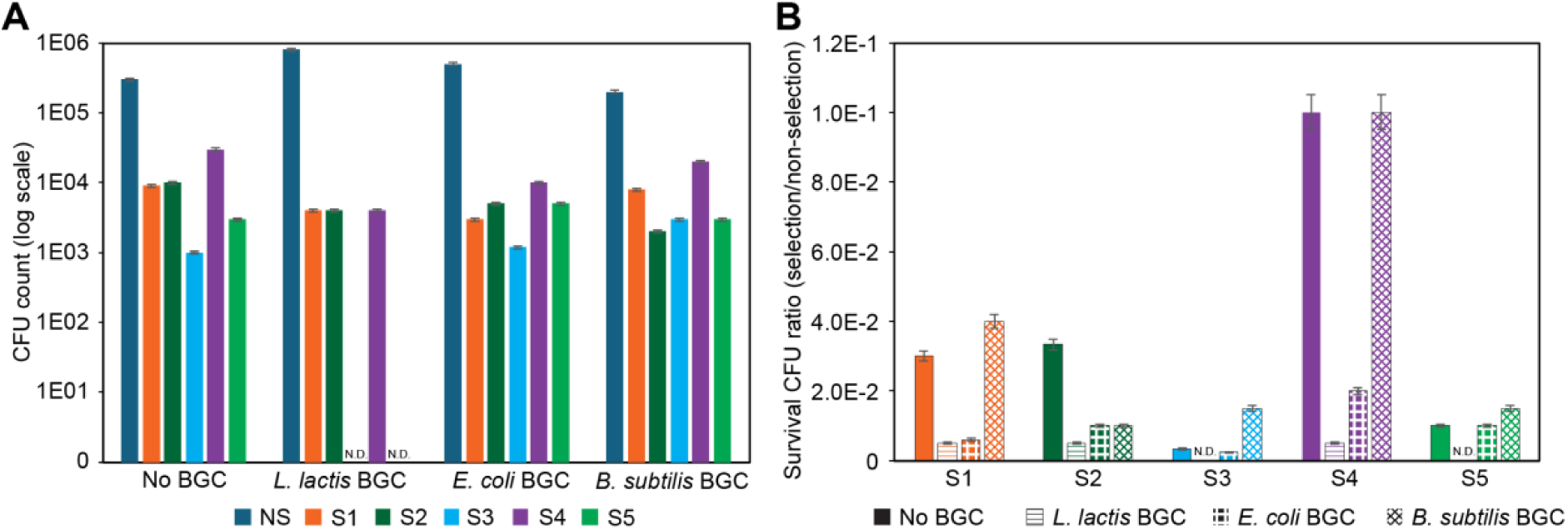
Survival colony counts in selective media after library transformation. (A) CFU count of library transformants in non-selective media (NS) and selective media (S1-S5) (B) CFU survival ratio of selective CFU count over CFU count in non-selective media (NS).

**Supplementary figure 5:**
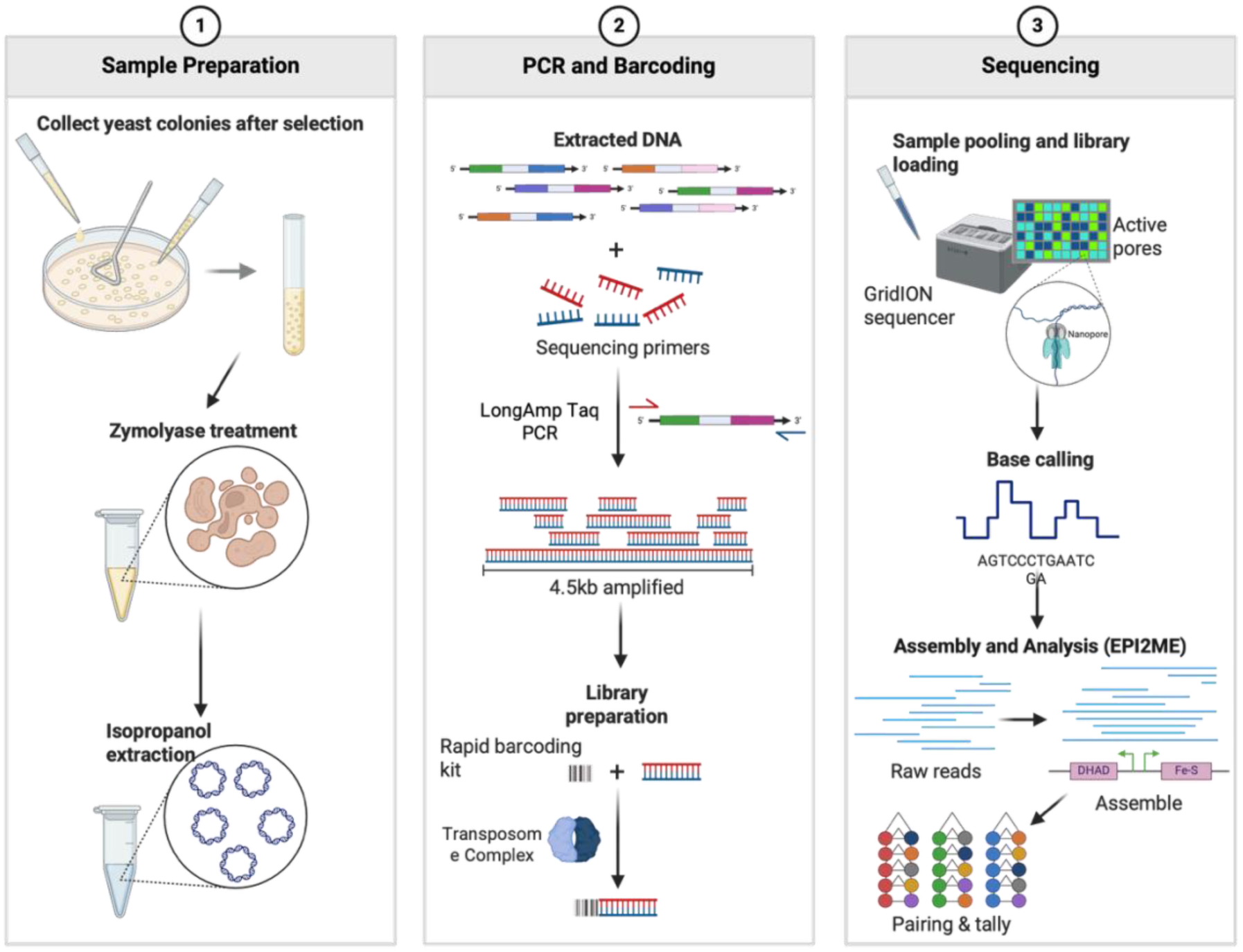
Framework of sequence processing. Sample preparation; colonies grown were collected in PBS buffer and cells were lysed with zymolyase. Isopropanol extraction was used to precipitate plasmid DNA. PCR and barcoding; extracted DNA was targeted by primers that flank outside of dual GAL promoter gene cassette and amplified by longAMP taq PCR. Amplification of the largest theoretical cassette (4.5kb) in the library was achieved. Amplified product was prepped via Oxford barcoding kit utilizing a transposome complex. Sequencing; processed samples were placed in Oxford GridION nanopore sequencer. Active pores were monitored, and results were processed via base calling, assembled, and analyzed with custom Python code and the Nextflow: EPI2ME alignment workflow.

**Supplementary figure 6:**
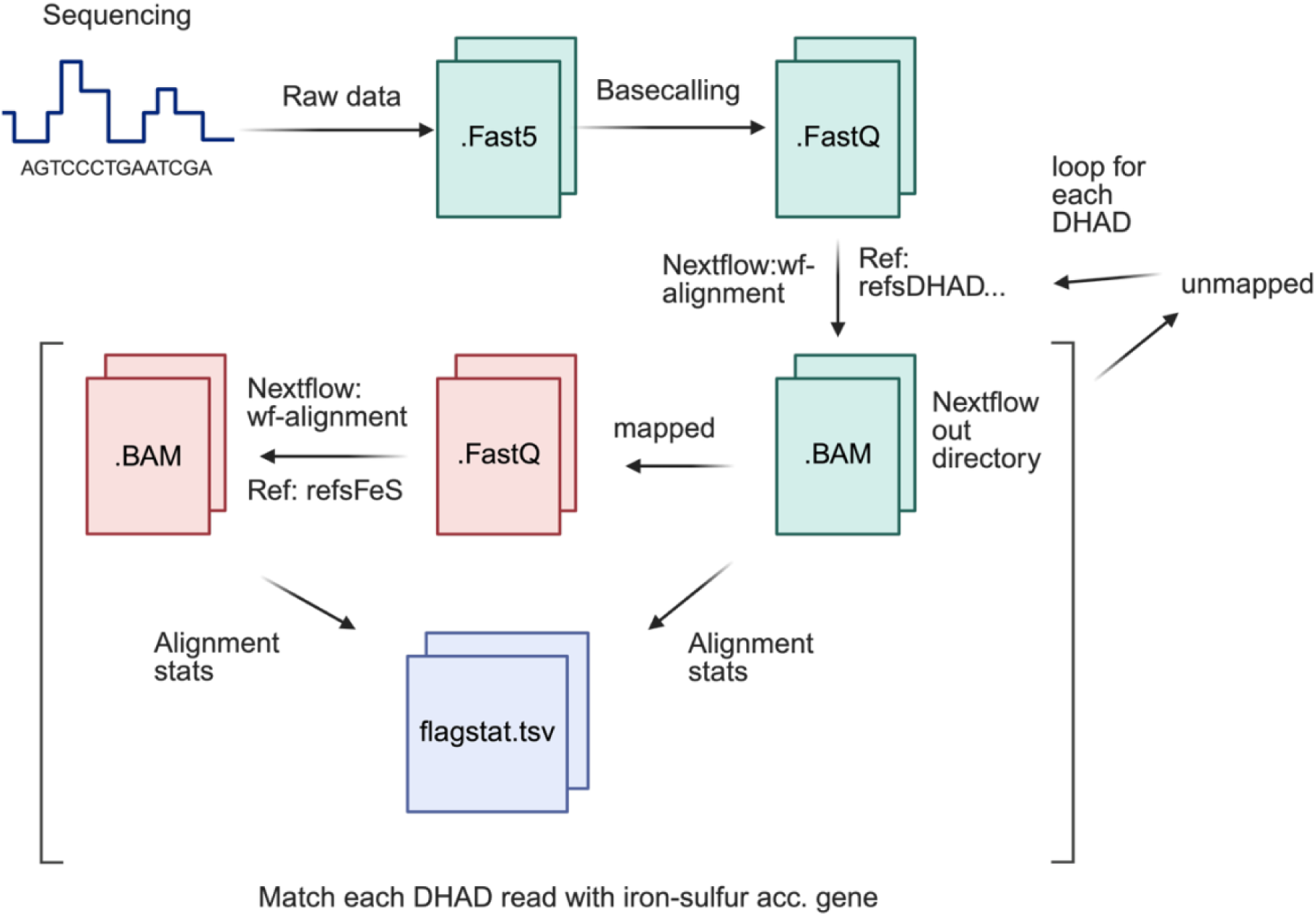
Sequence processing. Raw data from nanopore sequencing was converted to .FastQ format. Utilizing nextflow wf-alignment workflow (EPI2ME), reference DHAD orthologs were mapped and statistics were loaded into a .tsv file. Of the reads found to be mapped to a particular DHAD ortholog, they were mapped with reference iron-sulfur cluster accessory genes. Alignment statistics were loaded on the compiled .tsv file. This process was repeated for each DHAD ortholog.

**Supplementary figure 7:**
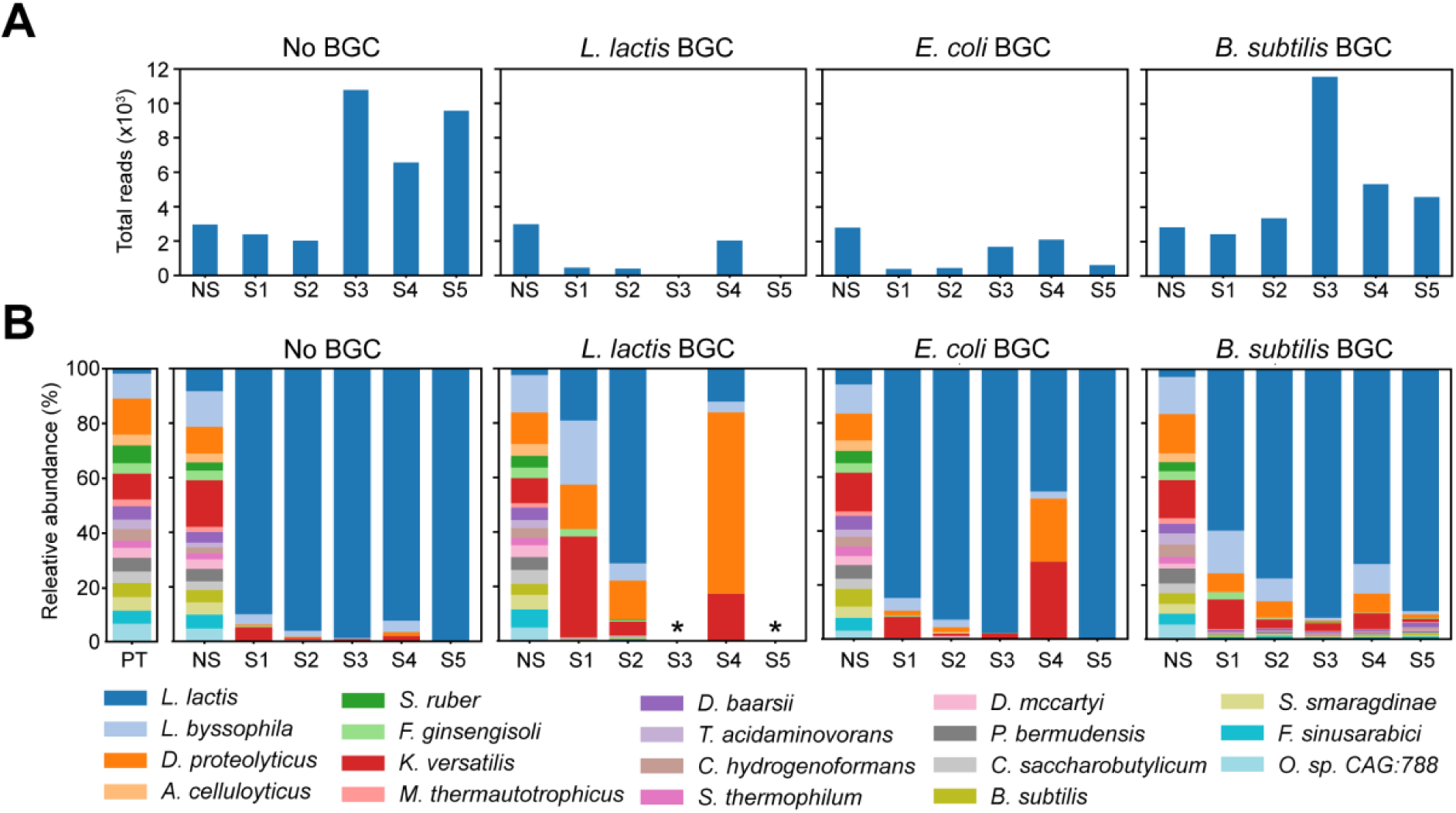
Raw reads, relative abundance (with L. lactis DHAD). (A) total reads of DHAD orthologs found in each media type from strains with *L. lactis*, *E. coli*, *B. subtilis,* or no BGC present. (B) Relative abundance of reads in each run. PT; high-copy plasmid library used to transform strains directly sequenced. * Indicates conditions with no observed colonies.

**Supplementary figure 8:**
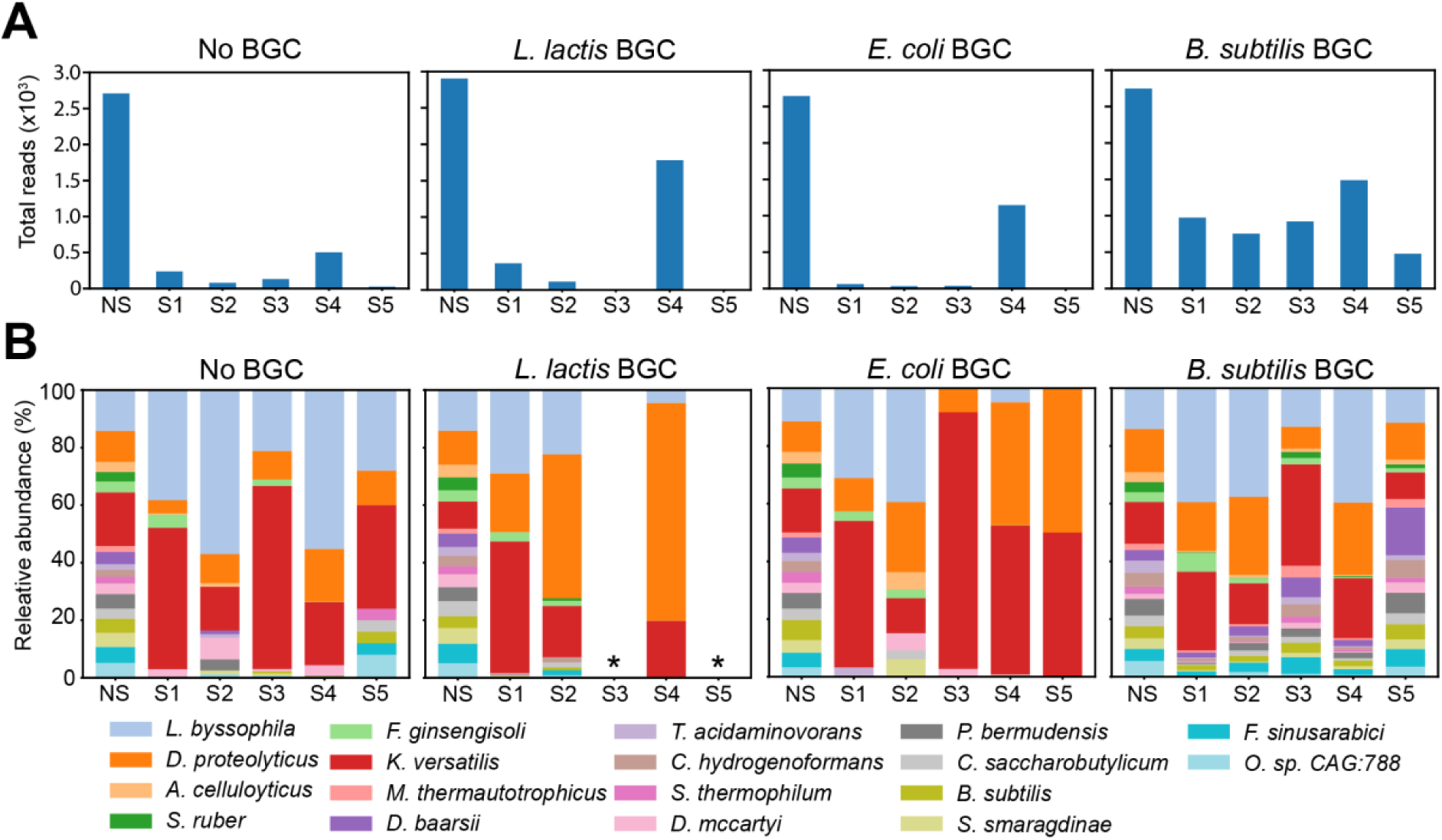
Raw reads, relative abundance (without L. lactis DHAD) (A) total reads of DHAD orthologs found in each media type from strains with *L. lactis*, *E. coli*, *B. subtilis,* or no BGC present. (B) Relative abundance of reads in each run. PT; high-copy plasmid library used to transform strains directly sequenced. * Indicates conditions with no observed colonies.

**Supplementary figure 9:**
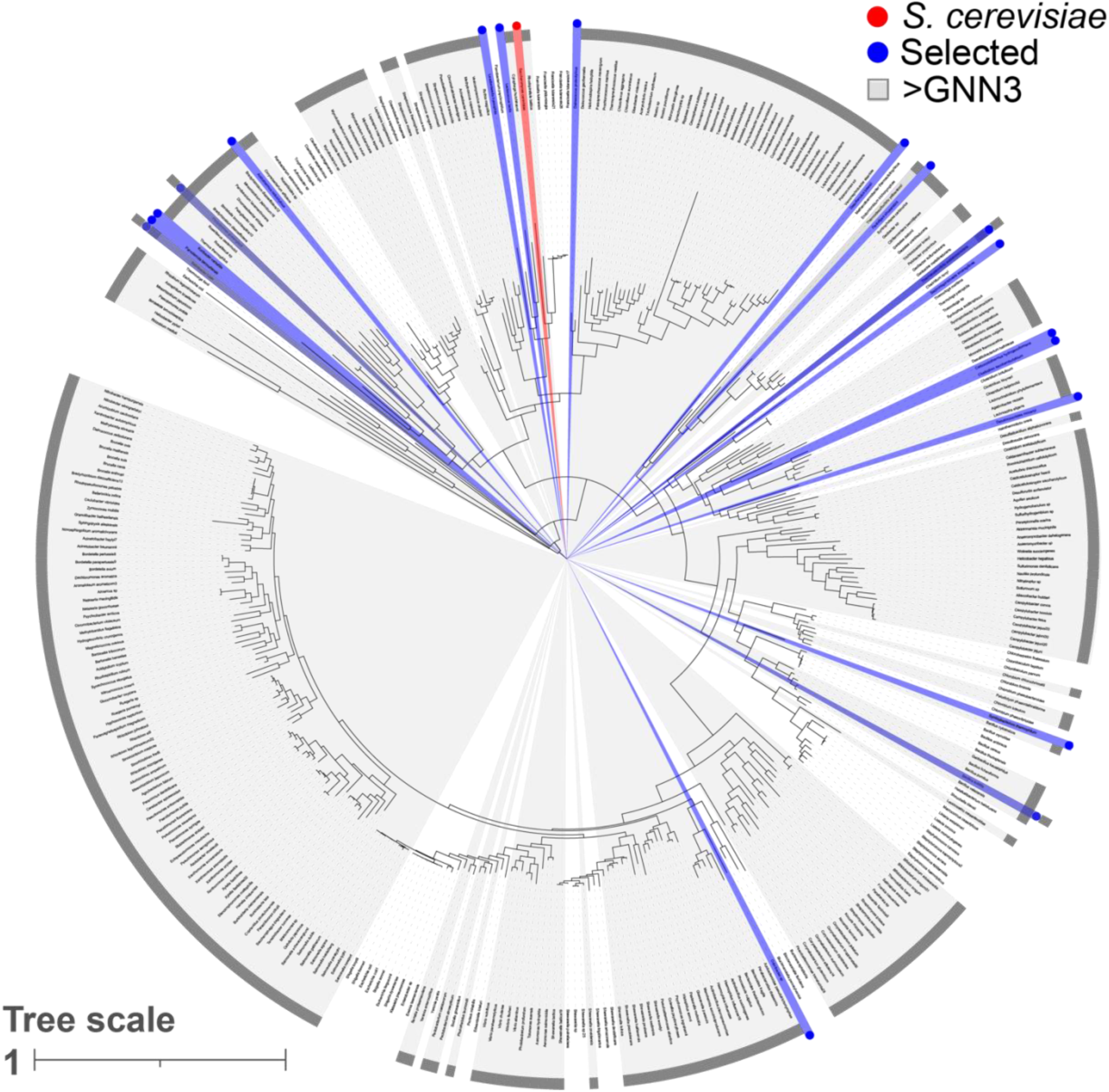
Figure 3A magnified.

**Supplementary figure 10:**
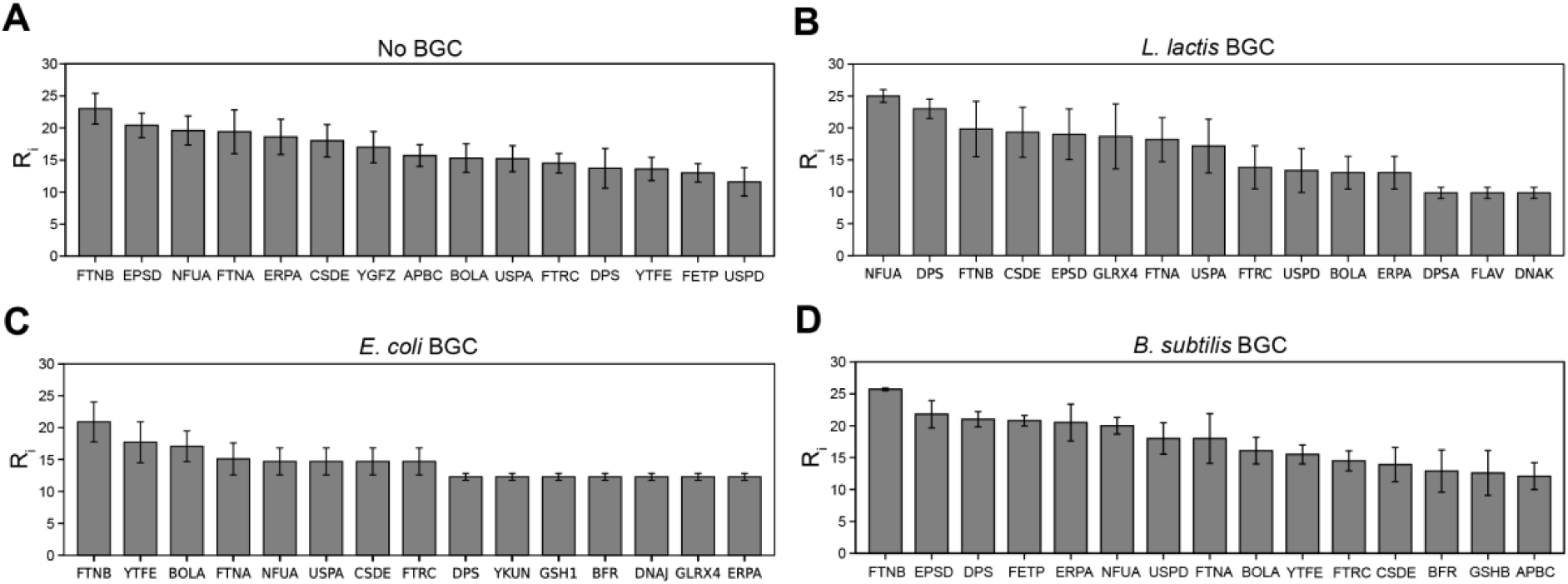
Ranking of most frequent iron-sulfur cluster accessory genes. Ranking scores of the most frequent iron-sulfur cluster accessory gene reads with DHAD orthologs when co-expressed with (A) no BGC, (B) *L. lactis* BGC, (C) *E. coli* BGC, and (D) *B. subtilis* BGC. Ri indicates ranking score; highest values are most frequent in the sample.

